# A TREX1 model reveals double-strand DNA preference and inter-protomer regulation

**DOI:** 10.1101/2022.02.25.481063

**Authors:** Wayne O. Hemphill, Thomas Hollis, Freddie R. Salsbury, Fred W. Perrino

## Abstract

The TREX1 3’ → 5’ exonuclease degrades DNA *in vivo* to prevent chronic immune activation through the cGAS-STING pathway. TREX1 degrades ss- and dsDNA containing a free 3’-hydroxyl, but the precise nature of immune-activating DNA remains an open question. The TREX1 homodimer structure is critical for exonuclease activity with amino acids from one protomer acting across the dimer interface contributing to catalysis in the opposing protomer. The unique TREX1 obligate homodimer structure suggests an intricate connection between the TREX1 protomers that has yet to be explained. We used biochemical assays, molecular dynamics simulations, and kinetic modeling to determine relative TREX1 affinities for ss- and dsDNA and to interrogate inter-protomer communication within the TREX1 homodimer. These new findings indicate that TREX1 is a semi-processive exonuclease with at least a 20-fold greater affinity for dsDNA than for ssDNA. Furthermore, we find extensively correlated dynamics between TREX1 protomers revealing newly identified substrate interactions in the TREX1 enzyme. These data indicate that TREX1 has evolved as a semi-processive exonuclease with a likely *in vivo* function to degrade dsDNA, where the TREX1 homodimer structure facilitates a mechanism for efficient binding and catabolism of dsDNA. These studies identify previously unrecognized regions of the TREX1 enzyme involved in DNA interactions, and our findings contribute to an emerging model of TREX1 exonuclease activity.

## Introduction

Three-prime repair exonuclease 1 (TREX1) degrades DNA in the 3’ → 5’ direction to prevent aberrant DNA-sensing by cyclic GMP-AMP synthase (cGAS) and subsequent activation of the stimulator of interferon genes (STING) pathway (1, 2). TREX1 mutations are associated with a spectrum of autoimmune disorders including Aicardi-Goutieres syndrome (AGS), systemic lupus erythematosus (SLE), familial chilblain lupus (FCL), and retinal vasculopathy with cerebral leukodystrophy (RVCL) (2). TREX1 exhibits robust exonuclease activity *in vitro* with ss- and dsDNA containing a free three-prime hydroxyl (3–6). However, cGAS is the principle DNA sensor driving pathology in *TREX1* mutant mice (7–9), and cGAS activation requires dsDNA binding (10). Furthermore, the *TREX1* p.D18N, p.D18H, p.D200N, and p.D200H mutations exhibit dominant disease genetics *in vivo* (2, 11), and the recombinant TREX1^D18N^, TREX1^D18H^, TREX1^D200N^, and TREX1^D200H^ enzymes recapitulate these dominant biochemical properties during dsDNA degradation of plasmid and chromatin DNA *in vitro* (11–13). These findings support the concept that failure to degrade dsDNA is the principal pathway of immune activation in TREX1-mediated autoimmune disease.

TREX1 has an obligate homodimer structure with residues in one protomer providing necessary structural and catalytic elements that contribute to full catalytic activity during DNA degradation in the opposing protomer (4, 14–16). For example, the Arg-62 and Arg-114 residues reach across the TREX1 dimer interface directly contributing to catalysis in the opposing protomer providing a biological rationale for the TREX1 dimer structure. The unique TREX1 homodimer is kinetically stable and does not dissociate and re-equilibrate at measurable rates. This structural requirement for full catalytic competency suggests that the presence and conformation of DNA in one protomer might affect activity in the opposing protomer (15, 17). Inter-protomer regulation of catalysis is supported by the exonuclease activities of TREX1 heterodimers of dominant disease alleles (11, 13), which demonstrate that the D18N, D18H, D200N, and D200H heterodimers do not degrade dsDNA but degrade ssDNA at ~50% the rate of the TREX1 wild-type enzyme.

In this work we used biochemical exonuclease assays and kinetic modeling to determine the TREX1 binding and catalytic rate constants for ss- and dsDNA, and thermal shift assays to validate the implicated binding affinities. The TREX1 residues contributing to ss- and dsDNA binding were also interrogated with all-atoms molecular dynamics (MD) simulations. In addition, we coupled biochemical measurements of the exonuclease activities of TREX1 dominant mutant enzymes with MD simulations to interrogate potential mechanisms of inter-protomer communication in the TREX1 homodimer. Our findings reveal a dramatic TREX1 preference *in vitro* for dsDNA, and a mechanism for inter-protomer regulation of TREX1 dsDNA exonuclease activity, which collectively support previous findings that dsDNA is a critical TREX1 biological substrate.

## Results

### TREX1 thermostability is greater upon dsDNA binding compared to ssDNA

To assess TREX1 binding preferences between ss- and dsDNA, we compared thermostability effects of ss- vs dsDNA binding to human TREX1_1-242_ (hT1) and mouse TREX1_1-242_ (mT1). Ligand binding increases protein thermostability in a manner positively correlated to protein-ligand binding affinity (18). We used thermal shift assays (TSA) to compare the melting temperature (T_m_) of TREX1 in the absence of ligand and in the presence of ss- and dsDNA (Figure 1). The dsDNA binding conferred a much greater thermostability to the hT1 (ΔT_m_ ≈ +28.3 °C) compared to ssDNA binding (ΔT_m_ ≈ +13.6 °C), and similar results were obtained using mT1 (ΔT_m_ ≈ +20.1 vs +7.1 °C). These thermostability data indicate that TREX1 binds to dsDNA with greater affinity than ssDNA. We also note that the hT1 apoenzyme is considerably less thermostable compared to the mT1 (T_m_ = 51.5 °C vs 58.2 °C). In contrast, the hT1 and mT1 ssDNA and dsDNA complexes exhibit comparable thermostabilities (Figure 1). The increased thermostability of the hT1 and mT1 enzymes, most dramatically apparent in the presence of dsDNA, is a strong indication of dsDNA as the biochemically preferred ligand.

**Figure 1:**
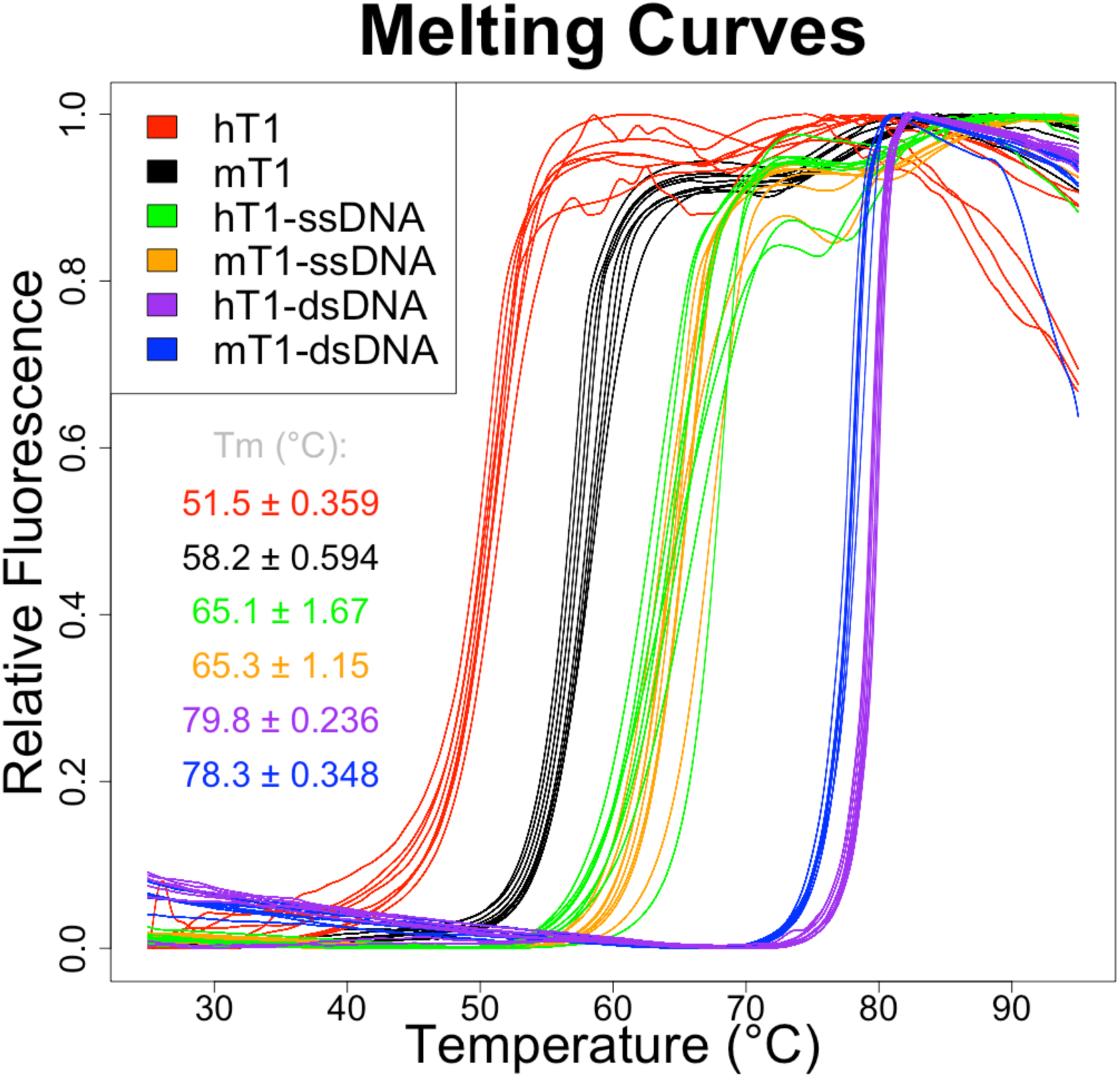
TREX1 binds dsDNA with greater affinity than ssDNA. The indicated TREX1 apoenzymes and TREX1-DNA complexes were subjected to an increasing temperature gradient in the presence of SYPRO Orange. Fluorescence was monitored as a metric of protein unfolding. Data shown is from a single experiment with 8 replicate reactions per condition. Graph was prepared in R v3.6.1. The temperatures associated with the inflection points of the curves were calculated as a measure of melting temperature. Values provided indicate mean ± standard deviation of melting temperature (Tm) across the 8 replicate reactions per condition. Calculations were performed in R v3.6.1, and graphic was prepared in PowerPoint (Microsoft).

### TREX1 steady-state kinetic constants indicate preference for dsDNA compared to ssDNA

TREX1 steady-state kinetic constants (K_m_ and k_cat_) were measured using ss- and dsDNA substrates to reveal any ligand preference (Figure 2 and Table 1). To quantify dsDNA degradation, we developed a novel absorbance-based DNA degradation assay using a singly nicked plasmid DNA (see Experimental Procedures). We first determined assay sensitivity by measuring known concentrations of dAMP to establish a standard curve (Figure 2A), and then determined the linear range of the assay in a time course reaction (Figure 2B). Initial velocities of hT1 degradation of the nicked-dsDNA plasmid were quantified in reactions using a range of substrate concentrations to generate a Michaelis-Menten curve (Figure 2C). These data indicate a k_cat_ ≈ 10 s^−1^ and K_m_ ≈ 10 nM for hT1 degradation of dsDNA. To quantify hT1 activity using ssDNA, we employed our previously established gel-based assay (19) using low (15 nM) and high (515 nM) ssDNA concentrations at three different hT1 concentrations (Figure 2D). The hT1 ssDNA k_cat_ ≈ 16 s^−1^ and K_m_ ≈ 120 nM for ssDNA were approximated from these ssDNA reactions and the derived equations (see Experimental Procedures). These steady state kinetic data affirm that hT1 has a greater binding affinity to dsDNA compared to ssDNA.

**Figure 2:**
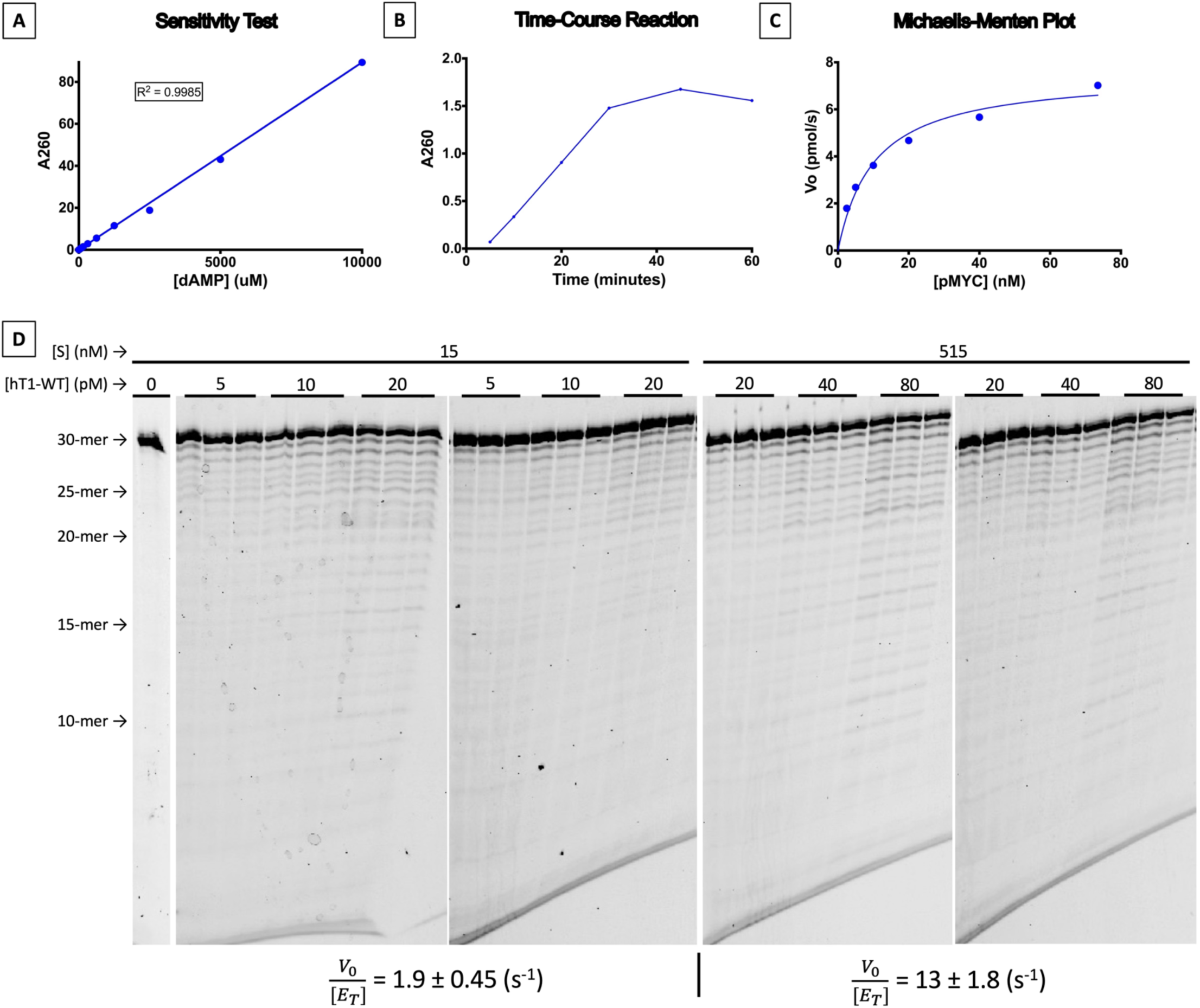
Determination of TREX1 dsDNA k_cat_ and K_m_. **[A]** *Test of absorbance assay sensitivity*. A_260_ was measured via NanoDrop for dNMP concentrations from 0.15 µM – 10 mM. Lower limit of nucleotide detection was ~5 µM, and the assay was linear to A_260_ ≥ 89. **[B]** *Test reaction with absorbance assay*. Time-course reaction was performed as described with standard reaction components, 30 nM hT1, and 60 ng/µL nicked-plasmid. **[C]** *Determination of TREX1 dsDNA kinetic constants*. Reactions containing standard components, 30 nM hT1, and the indicated concentrations of nicked dsDNA were incubated 10 minutes at ~21°C, quenched with EDTA, filtered, and their A_260_ measured via NanoDrop. Standard curve in panel A was used to calculate dNMP concentrations used for initial velocity calculations. Data represents a single experiment with single reactions for each substrate concentration. From this data it was calculated that k_cat_ = 10 s^−1^ and K_m_ = 10 nM. **[D]** *Exonuclease activities of TREX1 on ssDNA.* Standard exonuclease reactions were prepared with indicated concentrations of hT1, incubated at room temperature for 20 minutes, quenched in ethanol, then visualized on a 23% urea-polyacrylamide gel. Top bands indicate undegraded 30-mer oligonucleotide (’30-mer’), which TREX1 degrades in single-nucleotide units to generate the laddered bands (other ‘-mer’). Densitometric quantification of these gels was used to calculate activity rates. Values provided at the bottom of each gel set are mean and standard deviation. Activity rates are calculated for each substrate concentration from 18 total reactions each via 6 replicates per 3 concentrations of hT1. Figure was prepared in PowerPoint (Microsoft).

**Table 1:** Classic kinetic constants for TREX1. ‘Reported’ values are taken from prior work (4). dsDNA values from this study were calculated empirically using the described absorbance-based exonuclease assay. ssDNA values from this study were approximated via Monte-Carlo simulation using the ssDNA activity rates of wild-type enzyme that were calculated in Figure 2D. Values shown indicate *µ* ± *σ*, except for this study dsDNA values and the reported specific activities. ‘N/A’ = not applicable. ^†^ Concentration of occupied active-sties inferred to be half the protomer concentration; regression directly indicates k_cat_ ≈ 5.0 s^−1^. ^††^ K_m_^app^ ≈ [E_T_], it’s possible that K_m_ < K_m_^app^. Calculations and simulations were performed in R v3.6.1, and table was prepared in Word (Microsoft).

|  | ssDNA |  |  | dsDNA |  |  |
| --- | --- | --- | --- | --- | --- | --- |
| | $k_{cat}$ (s<sup>-1</sup>) | $K_m$ (nM) | $k_{cat} / K_m$ (M<sup>-1</sup>s<sup>-1</sup>) | $k_{cat}$ (s<sup>-1</sup>) | $K_m$ (nM) | $k_{cat} / K_m$ (M<sup>-1</sup>s<sup>-1</sup>) |
| *This Study* | 16 ± 3.2 | 120 ± 57 | 1.5 ± 0.42 (x10<sup>8</sup>) | <sup>†</sup>10 | <sup>††</sup>10 | 1.0 (x10<sup>9</sup>) |
| *Reported* | 22 ± 1.1 | 19 ± 4.0 | 1.2 (x10<sup>9</sup>) | 11 ± 0.77 | 15 ± 4.4 | 7.3 (x10<sup>8</sup>) |

### A TREX1 kinetic model supports higher binding affinity to dsDNA compared to ssDNA

A reaction scheme for TREX1 exonuclease activity was developed to estimate the TREX1 dsDNA and ssDNA rate constants (Figure 3 – scheme 1). Scheme 1 accounts for many established TREX1 properties including (1) the polymeric nature of TREX1 substrates (i.e., multiple excisable nucleotides in a single polynucleotide substrate molecule), (2) substrate concentration remaining constant between most TREX1 catalytic events (i.e., the number of 3’-hydroxyls on a polynucleotide is unchanged by nucleotide excision), (3) the potential for processive catalytic activity (i.e., multiple nucleotides excised from a polynucleotide between association and dissociation), (4) a substrate length-independent catalytic rate constant (i.e. comparable efficiency catabolism of polynucleotides with ≥4 nucleotides), and (5) irreversible catalysis (3–5, 20). This basic reaction scheme (Figure 3 – scheme 1) is expanded to describe competition between TREX1 variants (Figure 3 – scheme 2). Both reaction schemes 1 and 2 provide mathematical descriptions of reactions (Eq. 16.1–17.8) and the number of nucleotides processively excised by TREX1 (Eq. 18), which include TREX1 constants for off-rate (k_-1_), on-rate (k_1_), catalytic rate (k_2_), and processivity (Ψ).

**Figure 3:**
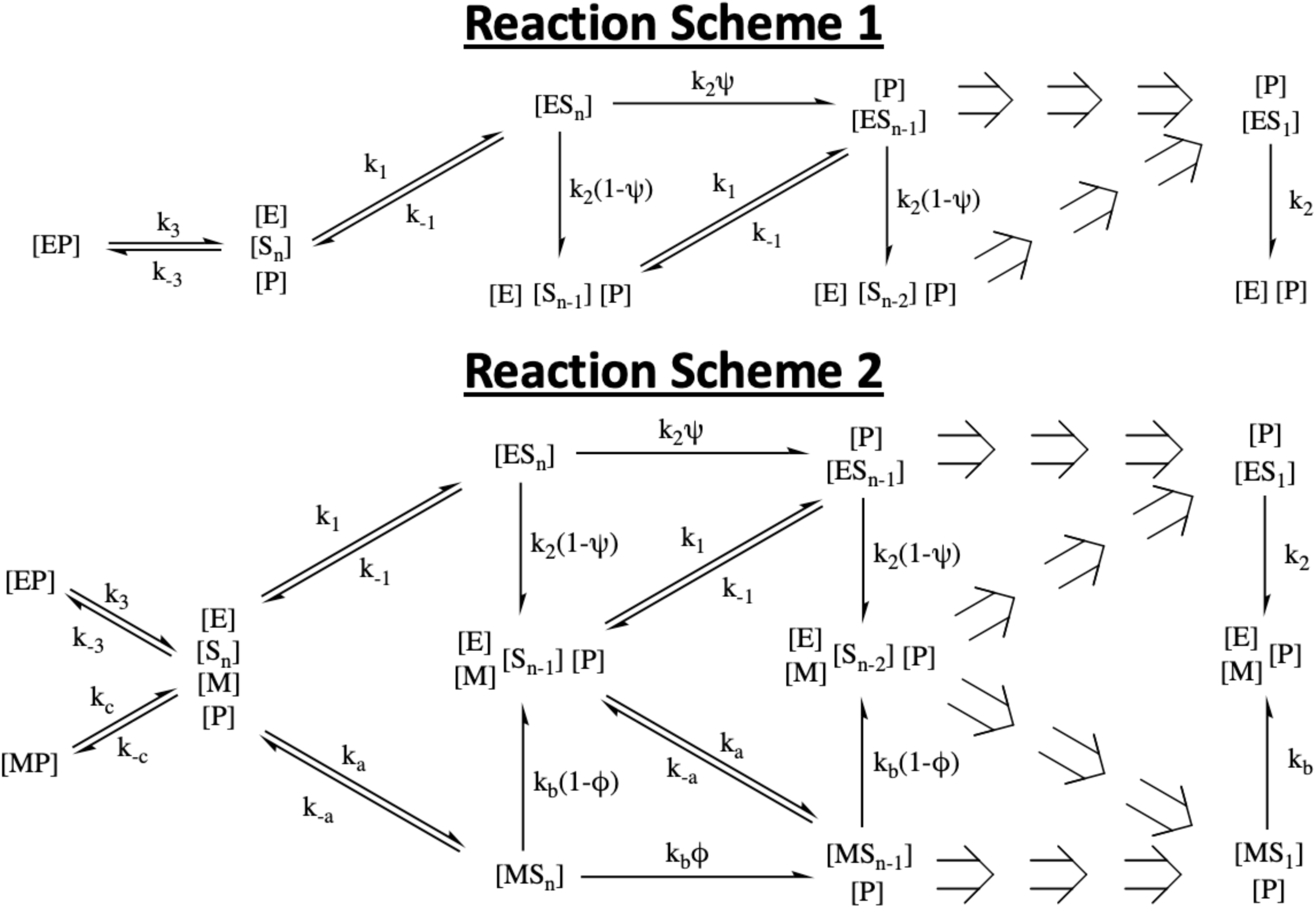
Proposed reaction schemes for TREX1 exonuclease activity and competition between TREX1 variants. Schemes are provided for exonuclease activity of wild-type TREX1 only (scheme 1), and for gross activity of TREX1 variants in competition (scheme 2). The schemes describe reactions composed of wild-type (E) and mutant (M) TREX1 enzyme, polynucleotides (S) with varying (n) excisable nucleotides (S_n_), and excised dNMPs (P), where conjugations of these reactants are complexes, and bracketed terms indicate concentrations. Transitions between reaction states are described by rate constants (wild-type, mutant) for association (k_1_, k_a_), dissociation (k_-1_, k_-a_), and catalysis (k_2_, k_b_), where catalytic rate constants are adjusted with a processivity parameter (Ψ, Φ).

We used data from our steady-state kinetic studies and prior studies to solve the TREX1 ssDNA rate constants in our reaction scheme (see Experimental Procedures, Supplemental 1 and Figure 4A). Our findings indicate that the TREX1 ssDNA equilibrium dissociation constant (K_d_) is 42 nM and Ψ is 0.75 representing a 25% chance that TREX1 dissociates upon excising a single nucleotide from ssDNA (Table 2). For dsDNA, insufficient data was available for a similar approach. However, prior studies indicate that the TREX1 D18N mutation abolishes catalysis without significant effects on binding affinity, and that recombinant human TREX1_1-242_^D18N/D18N^ (hT1-DN) enzyme competitively inhibits wild-type enzyme on ds- but not ssDNA (13, 17, 19, 21, 22). These data indicate that this substrate-specific inter-enzyme inhibition is a manifestation of different TREX1 ss- and dsDNA rate constants. We performed fluorescence-based assays (19) to measure the effects of hT1-DN competition on hT1 exonuclease activity (Supplemental 2), and we used these data to approximate the TREX1 dsDNA rate constants in our reaction scheme (see Experimental Procedures). Our findings indicate that the TREX1 dsDNA K_d_ is about 20-100 pM and Ψ is about 0.75-0.85 (Table 2). We note that simulating such competition experiments with our ssDNA rate constants predicts no inhibition of hT1 ssDNA exonuclease activity by hT1-DN (not shown), consistent with prior findings (13, 17, 19, 21, 22). Thus, our kinetic modeling from experimental data suggests that the TREX1 ds- versus ssDNA K_d_ is 400- to 2000-fold lower, consistent with the substantially greater TREX1 ds- versus ssDNA binding thermostabilities (Figure 1).

**Figure 4:**
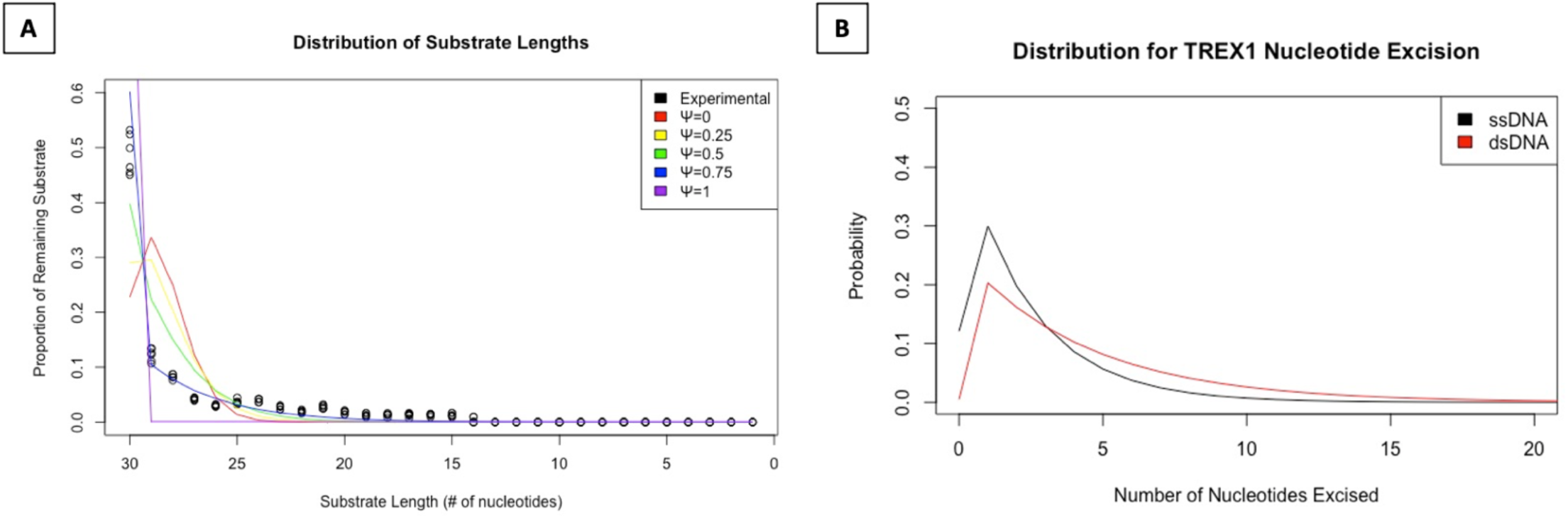
TREX1 ssDNA degradation is consistent with modest processivity. **[A]** A polyacrylamide gel-based TREX1 ssDNA degradation assay was performed as described in the relevant methods section, and the banding data quantified to give the proportion of total substrate concentration for each length substrate remaining. Substrate length proportions for all 6 replicate reactions are displayed (‘Experimental’). The five optimized sets of Ψ, k_1_, and k_-1_ parameter values identified in Supplemental 1 were used to simulate their predicted results for the experiment, which are indicated in the graph legend by their corresponding Ψ value. Simulated reactions are defined in Supplemental 6. **[B]** The TREX1 kinetic constants in Table 2 were used with Eq. 18 to generate the predicted probability distribution for the number of nucleotides TREX1 excises from dsDNA and ssDNA substrates after initial enzyme-substrate complexing. For dsDNA, the matched kinetic constant values associated with Ψ = 0.80 were used. Calculations and plotting were performed in R v3.6.1.

**Table 2:** Kinetic constant approximations for TREX1 exonuclease activity model. A table of the values for relevant kinetic constants. Bracketed values are approximation pairs, where the first values in each bracketed vector are true under the same assumptions as one another. Other values are broadly applicable approximations. ‘n.d.’ is not determined. ^†^ Values taken from Table 1, which were constrained in the modeling. Calculations were performed in R v3.6.1, and table was prepared in Word (Microsoft).

| | $k_{-1}$ (s<sup>-1</sup>) | $k_1$ (M<sup>-1</sup>s<sup>-1</sup>) | $k_2$ (s<sup>-1</sup>) | $k_{-3}$ (s<sup>-1</sup>) | $k_3$ (M<sup>-1</sup>s<sup>-1</sup>) | $\psi$ |
| --- | --- | --- | --- | --- | --- | --- |
| *ssDNA* | 2.2 | 5.2 (x10<sup>7</sup>) | <sup>†</sup>16 | n.d. | n.d. | 0.75 |
| *dsDNA* | ~ [0.10,0.050,0.020] | ~1.0 (x10<sup>9</sup>) | <sup>†</sup>10 |  |  | ~ [0.75,0.80,0.85] |

Finally, we used the inferred TREX1 rate constants (Table 2) to predict (via Eq. 18) the number of nucleotides TREX1 is likely to excise processively on ss- versus dsDNA substrates. Our findings indicate that processive excision of more than 10 nucleotides from either substrate is unlikely (Figure 4B). Notably, while processive excision of at least 7 nucleotides is more likely for ds- than ssDNA substrates, the absolute probability of such events is still low. Thus, TREX1 is predicted to excise short stretches of nucleotides from DNA following association events.

### MD simulations implicate differential TREX1 binding dynamics for ss- vs dsDNA

Our above data indicate significantly different TREX1 binding affinities for ss- versus dsDNA, which suggests differential protein-polynucleotide interactions between the two substrates. Crystal structures reveal static protein-polynucleotide contacts that are highly conserved between the two substrates (16, 23). To better define the transient protein-polynucleotide interactions that likely contribute to the differential TREX1 binding affinities for ss- versus dsDNA, we performed all-atoms molecular dynamics (MD) simulations with mT1 as apoenzyme, bound to 4-mer oligo, and bound to dsDNA.

Root mean squared deviation (RMSD) comparisons of mT1 against initial conditions shows that all simulations quickly equilibrate without indications of instability (Supplemental 3). Visualization of the conformational free energy maps of the mT1 backbone alpha carbons (Cα) in each of these systems reveals that mT1 conformations are significantly altered by its ligand binding state, and that ss- versus dsDNA binding have differential effects on mT1 conformation (Supplemental 4). Root mean squared fluctuation (RMSF) analysis of the Cα in each system identified three areas of differential fluctuation in the mT1 backbone, which have the potential for direct interaction with substrate in the active site (Figure 5A). Notably, the TREX1 flexible loop (aa. 164-175) seems differentially impacted by the two substrates, being significantly rigidified during dsDNA occupation of the active site (Figure 5A – green markers). We also identified a short region (aa. 23-27) with the capacity to contact the 3’-terminal nucleotides of each substrate, which is mildly rigidified during 4-mer occupation and further rigidified by dsDNA occupation (Figure 5A – purple markers). Another region (aa. 75-88) also exhibited greater rigidity with dsDNA bound, but no significant change in rigidity during 4-mer occupation (Figure 5A – orange markers). These simulations identify previously unrecognized regions of TREX1-DNA interactions.

**Figure 5:**
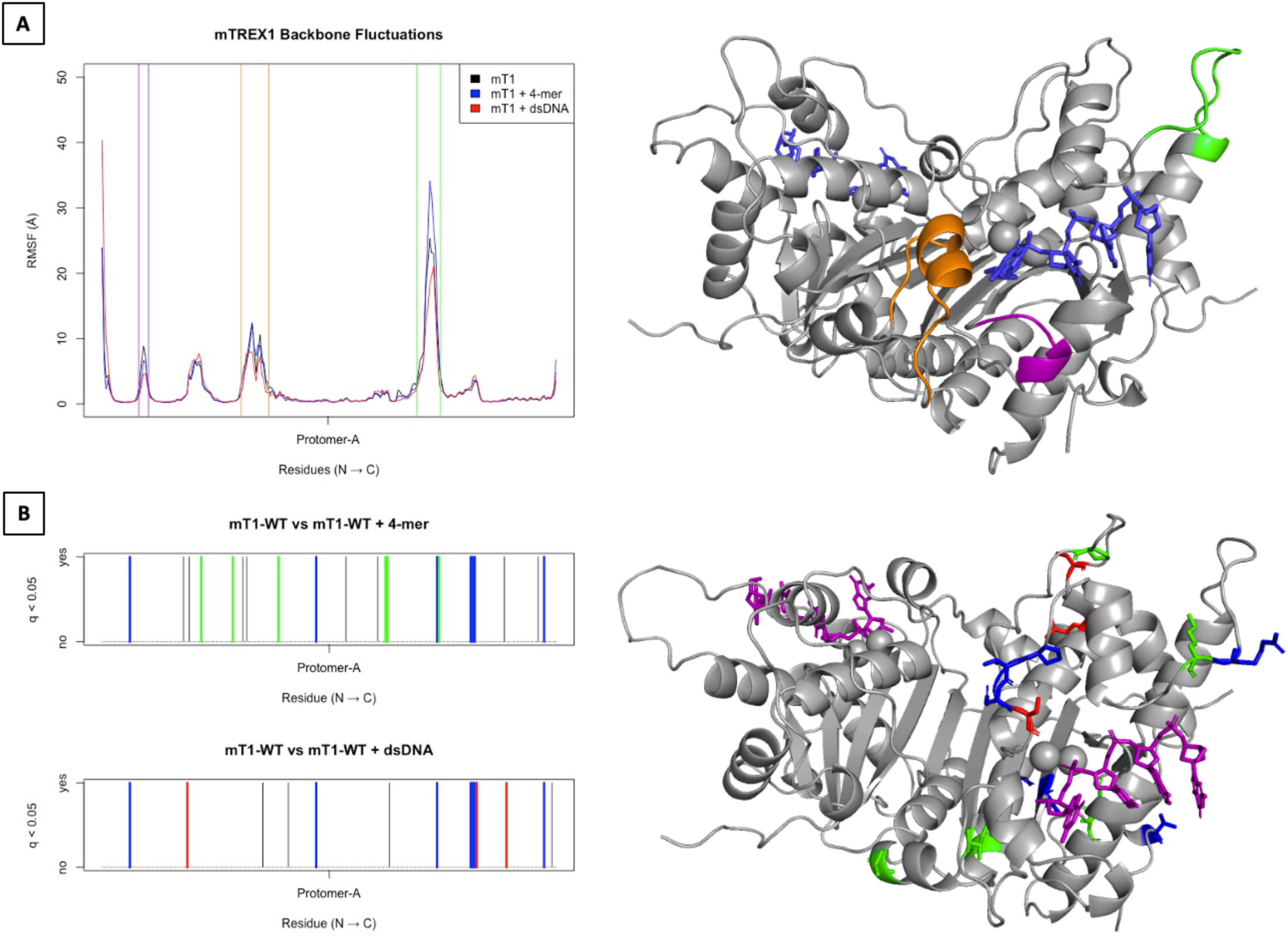
Active site occupation by ssDNA versus dsDNA is associated with differential mTREX1 dynamics. **[A]** *Comparative RMSF plots of TREX1 backbone fluctuation*. All TREX1 Cα atoms were used in an RMSF calculation, as described in the relevant methods section. Plot lines displayed are mean RMSF values from the four simulations per system. The purple, orange, and green vertical lines define regions of interest, which are similarly colored in panel E. All calculations and plotting were performed in R v3.6.1. A mT1:4-mer structure modeled from a mT1 crystal structure (PDB = ‘2IOC’) to include disordered flexible loops is included for reference. Both TREX1 protomers and all magnesium ions are colored grey, both ssDNA polymers are displayed as blue sticks. The dynamic regions identified in the RMSF plot are colored correspondingly. Graphic was constructed in PyMOL (36). **[B]** *Barcode graphs of residues with significantly different phi-psi angles*. For each residue, hierarchal clustering was used to determine conformations across all time frames, simulations, and systems. Residues were tested for differing conformation proportions between specific systems via chi-square tests followed by false discovery rate adjustment (q-values). Further details are found in the relevant methods section. Barcode plots show residues with significantly different conformation proportions between systems, with the title indicating the systems compared. Residues with blue markers had their conformations significantly altered by the presence of 4-mer and dsDNA in the active site. Green markers indicate residues whose conformations were only significantly altered by the presence of 4-mer in the active site. Red markers indicate residues whose conformations were only significantly altered by the presence of dsDNA in the active site. Analyses and graphing were performed in R v3.6.1. A mT1:4-mer structure modeled from a mT1 crystal structure (PDB = ‘2IOC’) to include disordered flexible loops is included for reference. Both TREX1 protomers and all magnesium ions are colored grey, both ssDNA polymers are displayed as purple sticks. The residues identified in the barcode graphs by colored markers are shown as correspondingly colored sticks. Graphic was constructed in PyMOL (36).

We subjected the amino acid phi/psi angles across all simulations and systems to a clustering analysis. Pairwise statistical comparisons of residue conformations were made between the systems and adjusted for false discovery rate (Figure 5B). As expected, several amino acid conformations were significantly altered during both 4-mer and dsDNA occupation of the active site (Figure 5B – blue markers). These included R174, L19, and aa. 191-193, all of which are adjacent to the active site with potential for DNA interactions. For several residues including S48, S194, and Q209, conformation was only altered by dsDNA occupation (Figure 5B – red markers). Of these, S194 is closest to the active site, and adjacent to the H195 residue previously identified as a residue of interest in TREX1 crystal structures (12, 16) and patient data (2). Curiously, there were several residues with conformations that were affected exclusively during 4-mer occupation of the active site, but only K175 is in proximity to the active site (Figure 5B – green markers). Collectively, these data affirm differential TREX1-DNA interactions for ds- versus ssDNA, implicate previously unrecognized TREX1 structural regions (aa. 23-27 and 75-88) as DNA binding contributors, and confirm prior implications that TREX1 flexible loop primarily binds dsDNA (13).

### Kinetics of dominant TREX1 mutants suggest inter-protomer activity regulation

The TREX1 dominant mutant heterodimers (e.g., D18N, D200N, etc.) exhibit exonuclease activity on nicked plasmid dsDNA comparable to that of the mutant homodimers, suggesting a mechanism for mutant protomer-mediated suppression of wild-type protomers in such a TREX1 heterodimer (11, 13, 19, 22). We interrogated this mechanism by quantifying exonuclease activities for the hT1, hT1-DN, and recombinant human TREX1_1-242_^WT/D18N^ (hT1-DNhet) enzymes at sub-enzyme concentrations of the nicked-plasmid substrate and at saturating concentrations of a 30-bp dsDNA (Figure 6A-B). We observed significantly greater exonuclease activities for hT1-DNhet and a 1:1 hT1:hT1-DN mix on the oligo versus plasmid dsDNA, consistent with the plasmid dsDNA sterically hindering simultaneous substrate degradation in both protomers of the TREX1 homodimer (Figure 6C). Consequently, our calculation of the TREX1 dsDNA catalytic rate constant was adjusted (Table 1). However, we note that the heterodimer was still ~2-fold less active on the oligo dsDNA than the 1:1 hT1:hT1-DN mix, despite these reactions having the same total concentrations of wild-type and D18N protomers, so some degree of regulated binding and/or catalysis across the TREX1 homodimer interface should not be ruled out.

**Figure 6:**
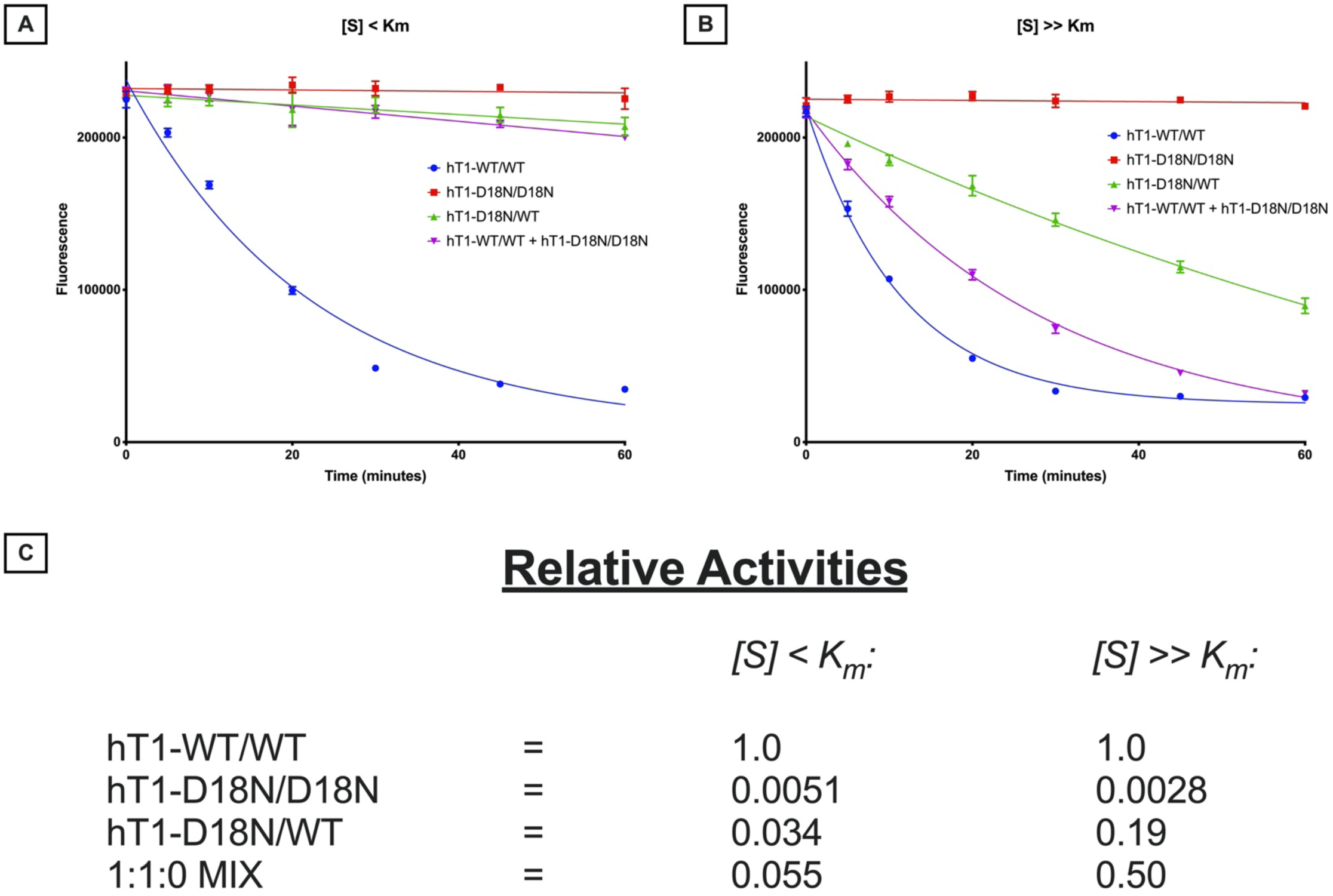
TREX1^D18N^ heterodimer deficiency in dsDNA exonuclease activity is dependent on substrate conditions. **[A-B]** *Fluorescence-Based Quantification of TREX1 dsDNA Degradation*. Standard exonuclease reactions were prepared with equimolar concentrations of the indicated enzymes, incubated at room temperature for the indicated times, quenched in SYBR Green, and dsDNA content measured by fluorescence. Plots of fluorescence vs time were generated and fit with one-phase decay nonlinear regression in Prism 9.0 (GraphPad). Plots are composites of 6 different reactions and representative of three experiments. Data points indicate mean, and error bars represent standard deviation. **[C]** *Activity Rates of TREX1 Variants*. Initial velocities were quantified from the respective regression lines in panels A-B and normalized to wild-type initial velocity to calculate relative activity. ‘[S] < K_m_’ refers to ~1 nM of a 10-kb dsDNA plasmid, and ‘[S] >> K_m_’ refers to ~1 µM of a 30-bp self-annealing oligo. For the 1:1:0 enzyme mix, mix ratios indicate the prevalence of hT1^+/+^, hT1^MUT/MUT^, and hT1^MUT/+^, respectively. All reactions have equimolar concentrations of total hT1, even mixes.

### MD simulations implicate inter-protomer communication between specific TREX1 regions

Our dominant TREX1 mutant kinetics experiments raise the question whether dsDNA binding in one active site could be communicated across the TREX1 dimer interface. To interrogate this possibility, we performed all-atoms MD simulations with mT1 apoenzyme and mT1 bound to dsDNA in a single active site (‘Protomer-A’) to compare dynamics of the unbound protomer (‘Protomer-B’) between systems. RMSD comparisons of mT1 against initial conditions shows that all simulations quickly equilibrate without instability (Supplemental 3). Visualization of the conformational free energy maps for the mT1 backbones reveals differential unbound protomer dynamics between these systems (Supplemental 5). An RMSF analysis revealed three distinct regions of the mT1 backbone that were rigidified by dsDNA occupation of the opposing protomer active site (Figure 7A). These include the TREX1 flexible loop (aa. 164-175), whose corresponding region in the related three-prime repair exonuclease 2 (TREX2) enzyme has been previously implicated in DNA-binding (24), and two previously unrecognized regions surrounding the active site (aa. 23-27 & 75-88). A phi/psi angle clustering analysis to identify conformationally altered residues echoed these findings, with most identified residues being located on the TREX1 flexible loop (Figure 7B). Importantly, these are the same amino acids impacted by DNA occupation in the resident protomer (Figure 5A).

**Figure 7:**
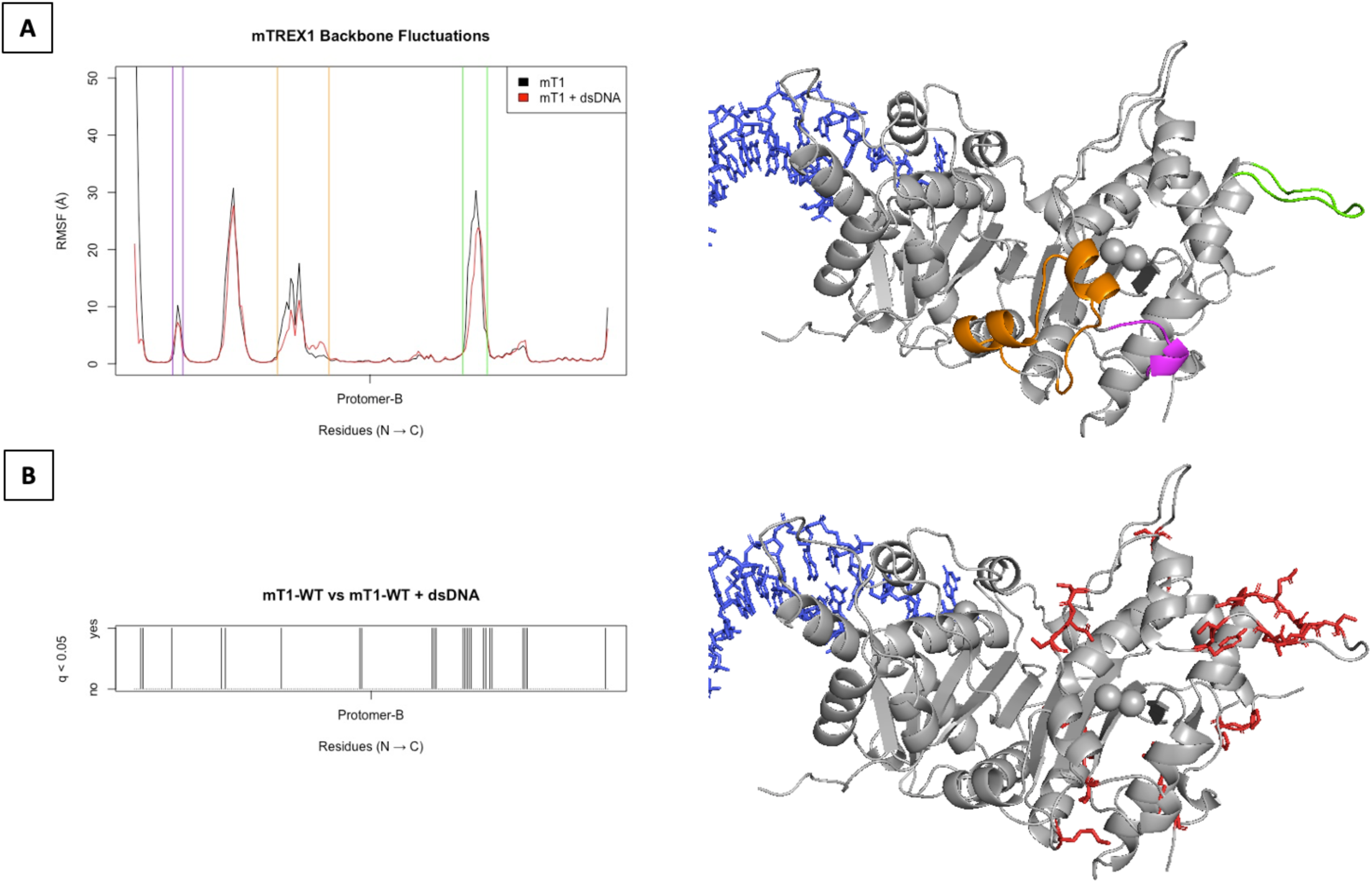
Active site occupation by dsDNA is associated with differential dynamics in the opposing protomer. **[A]** *Comparative RMSF Plots of mTREX1 Backbone*. All TREX1 Cα atoms in the unoccupied protomer of each system (‘Protomer-B’) were used in an RMSF calculation, as described in the relevant methods section. Plot lines displayed are mean RMSF values from the four simulations per system. The purple, orange, and green vertical lines define regions of interest, which are similarly colored in the reference mT1 crystal structure (PDB = ‘2IOC’) modeled to include disordered flexible loops. Both TREX1 protomers and all magnesium ions are colored grey, and the dsDNA polymers are displayed as blue sticks. Graphic was constructed in PyMOL (36). **[B]** *Barcode Graph of Residues with Significantly Different Phi-Psi Angles*. For each residue, hierarchal clustering was used to determine conformations across all time frames, simulations, and systems. Residues were tested for differing conformation proportions between specific systems via chi-square tests followed by false discovery rate adjustment (q-values). Further details are found in the relevant methods section. Barcode plots show residues with significantly different conformation proportions between systems, with the title indicating the systems compared. Analyses and graphing were performed in R v3.6.1. A mT1 structure modeled from a mT1 crystal structure (PDB = ‘2IOC’) to include disordered flexible loops is included for reference. Both TREX1 protomers and all magnesium ions are colored grey. The residues identified in the barcode graph are shown as red colored sticks. Graphic was constructed in PyMOL (36).

To interrogate general DNA-mediated inter-protomer regulation in the TREX1 homodimer, Cα correlation analyses were performed for mT1 and an mT1:4-mer complex (Figure 8). We found positively correlated dynamics between the protomer α5 helices (Figure 8 – correlation 1) and between the protomer β3 strands (Figure 8 – correlation 3), which were independent of 4-mer presence in the active sites. These observations are consistent with crystallographic studies that indicate extensive contacts at the TREX1 dimer interface to mediate structural stability (12, 15, 16). We also found inter-protomer dynamics correlations that were dependent on the presence of 4-mer in the active sites. These included positively correlated dynamics between the TREX1 flexible loop and α6 helix (Figure 8 – correlation 2) and between the flexible loop and aa. 79-93 (α3-α4) (Figure 8 – correlation 4), as well as negatively correlated dynamics between aa. 79-93 (α3-α4) and the TREX1 α5 helix (Figure 8 – correlation 2) and between aa. 79-93 (α3-α4) and aa. 22-31 (β1-α1-β2) (Figure 8 – correlation 6). These DNA-dependent correlations confirm prior studies indicating that the TREX1 flexible loop contributes to DNA binding (13, 24), and they implicate these additional regions in a DNA binding function.

**Figure 8:**
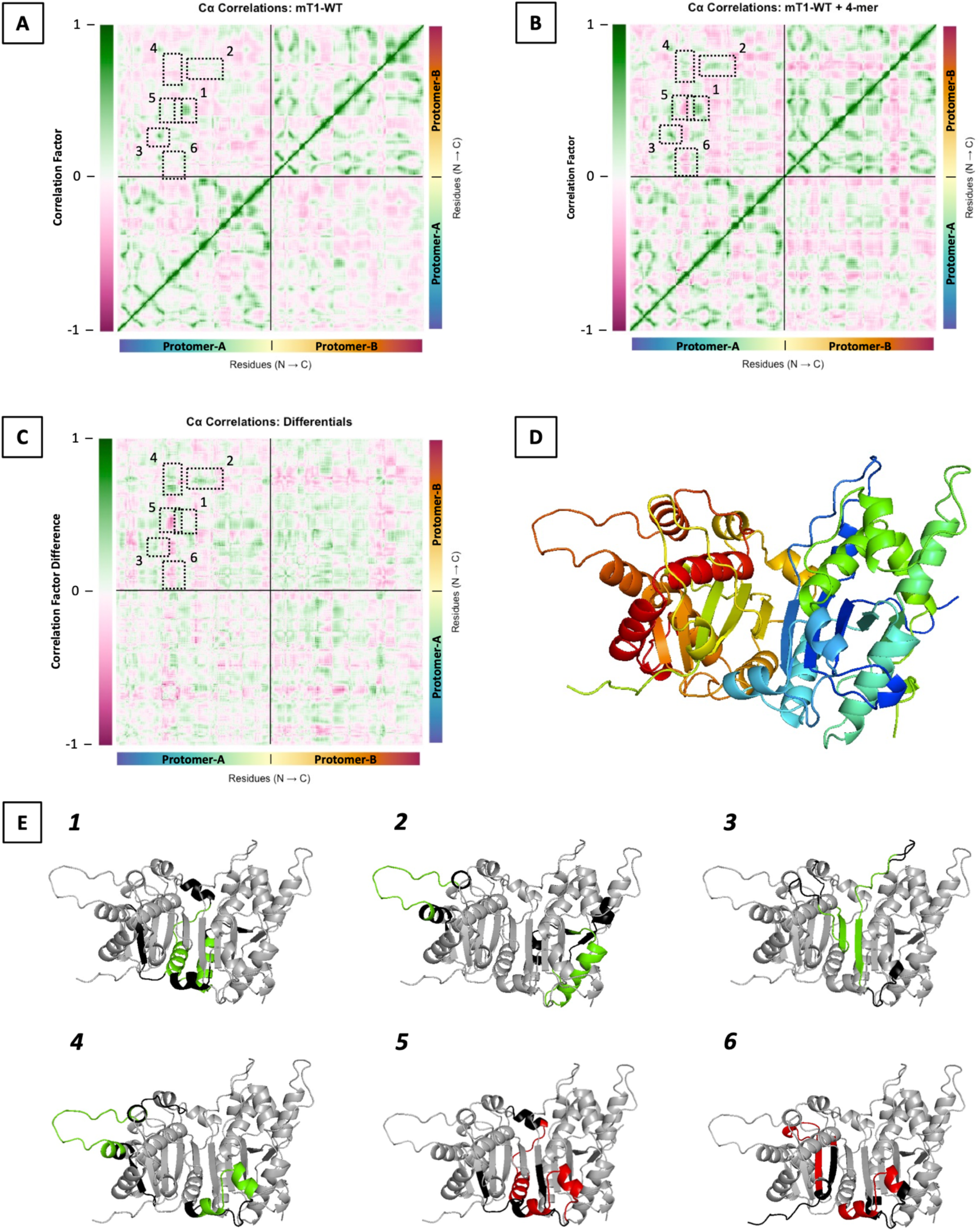
Regions of the mTREX1 structure associated with exonuclease activity have correlated dynamics between protomers. The α-carbons of every residue in the mT1 structure were subjected to correlation analyses against all other α-carbons in the mT1 structure, with comparisons across every time frame in every simulation, for each of the mT1 and mT1:4-mer systems. **[A-B]** *Correlation heat maps*. Bottom-x and right-y axes indicate the residues being compared, and the color gradient for these axis labels indicate the corresponding region of the mT1 structure shown in panel D. Vertical color legends to the left of each graph indicate the correlation factor between residues on the map, where ‘1’ is a perfect positive correlation of dynamics and ‘-1’ is a perfect negative correlation of dynamics. Solid black lines separate protomers, and dashed black boxes define regions of interest that are correspondingly numbered in panel E. Calculations and graphing were performed in R v3.6.1, and all other features were added in PowerPoint (Microsoft). **[C]** *Differential correlation heat map*. A heat map of the difference in correlation factors achieved by subtraction of panel A from panel B. All other features are as discussed for panels A-B. **[D]** *Color legend of TREX1 structure*. A mT1 structure modeled from a mT1 crystal structure (PDB = ‘2IOC’) to include disordered flexible loops. Backbone colors correspond to the bottom-x and right-y axes in panels A-C. Graphic was prepared in PyMOL (36). **[E]** *Structural representations of regions of interest*. mT1 structures modeled from a mT1 crystal structure (PDB = ‘2IOC’) to include disordered flexible loops. Italicized numbers indicate the correlated regions from panels A-C that are illustrated. The mT1 structures are generally shown as grey cartoons. The inner 50% of correlated areas from panels A-C are colored green or red for positively or negatively correlated regions, respectively. The peripheral 50% of correlated areas from panels A-C are colored black. Graphics were prepared in PyMOL (36).

## Discussion

Our kinetics and thermostability findings reveal a TREX1 biochemical substrate preference, collectively indicating a 400- to 2000-fold greater binding affinity for ds- versus ssDNA. We note that all prior reports (13, 15–17, 25) of TREX1 ssDNA activity rate (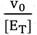) at a fixed substrate concentration (1.4-6.3 s^−1^ at 50 nM substrate) are consistent with our findings (Eq. 1 predicts ~4.7 s^−1^ at 50 nM substrate), with a modest deviation from the ssDNA K_m_ previously reported (4). Our MD simulations suggest that this difference in TREX1 ss- vs dsDNA binding affinity is largely attributable to interactions with the TREX1 flexible loop, as well as interactions with the TREX1 α1 (aa. 23-27) and α3 (aa. 75-88) helices. Mutation of the TREX1 flexible loop disproportionately affects ds- versus ssDNA exonuclease activity (13), the competitive inhibition of TREX1 wild-type enzyme by dominant mutants is significantly reduced upon co-mutation of the flexible loop residues (13), and the corresponding region in TREX2 contributes to DNA-binding (24). The TREX1 α1 (aa. 23-27) and α3 (aa. 75-88) helices are positioned near the 3’-terminal nucleotide binding pocket, indicating probable relevance to exonuclease activity (Figure 5A – orange and purple areas).

These findings provide insights into the likely nature of the TREX1 biological substrate and suggest an evolutionary preference for TREX1 degrading dsDNA species *in vivo*. TREX1 exhibits a comparable catalytic rate constant and high affinity for ssDNA, suggesting that TREX1 likely degrades ssDNA *in vivo*. Importantly, the 3’-terminal nucleotides of ss- and dsDNA substrates are positioned identically for catalysis in the TREX1 active site (16, 23), which explains the comparable TREX1 ss- and dsDNA catalytic rate constants. Similarly, sequence-independent contacts between TREX1 and the 3’-terminal nucleotides contributes to the affinity for ssDNA species. Thus, relative TREX1 binding affinity for ss- versus dsDNA seems the most appropriate metric for its substrate preferences, and this metric implicates dsDNA as the preferred TREX1 substrate.

Our kinetic data reveal additional features pointing to dsDNA as the preferred TREX1 biological substrate. Notably, the dsDNA on-rate is significantly higher than the ssDNA on-rate, approaching theoretical diffusion limits (26). These data are consistent with a mechanism where TREX1 “scans” the DNA backbone after initial association, allowing a more limited 1-dimensional search for free 3’-hydroxyls (24, 27–29). This scanning mechanism implicates a large biological substrate, where a scanning process is necessary to quickly identify 3’-hydroxyls and initiate degradation of potential immune-activating DNA polynucleotides. Our data do not address a possible ssDNA scanning mechanism since a much shorter ssDNA (~300-fold) was used in our experiments, so this proposed TREX1 scanning mechanism may be functional for both ssDNA and dsDNA. We note that assuming identical TREX1 ss- and dsDNA on-rate constants still predicts a 20- to 100-fold lower dsDNA K_d_. Despite the implication of a large dsDNA TREX1 substrate, our kinetic simulations suggest that the TREX1 catalytic cycle results in degradation of ≤ 10 nt of the scissile DNA strand. These TREX1 exonuclease properties parallel those of proofreading exonucleases (30), but loss of *TREX1* function does not result in a hypermutator phenotype (1, 2).

The TREX1 obligate homodimer structure is unique among exonucleases, suggesting a specific role during TREX1 degradation of its biological substrate (31, 32). Our studies of the TREX1 dominant mutant kinetics support substrate occupation of one TREX1 protomer being negatively associated with exonuclease activity in the opposing protomer. Further, our MD simulations suggest that the TREX1 flexible loop communicates across the dimer interface with α3-α4 (aa. 79-93) and with α6, and that α3-α4 (aa. 79-93) communicates across the dimer interface with β1-α1-β2 (aa. 22-31). This is consistent with TREX1 possessing flexible loop-dependent negative cooperativity across the dimer interface. Analogous studies in TREX2 also suggest that the flexible loop is involved in binding cooperativity (24, 27), and there is direct contact of the α6 and α3-α4 regions with the 3’-terminal nucleotide in existing crystal structures (16, 23). Our data suggest that the TREX1 α6 and/or α3-α4 regions are the ‘sensors’ of 3’-terminal nucleotide presence in the TREX1 active site and the flexible loop is the ‘effector’ of negative binding cooperativity.

Our findings support large dsDNA as the TREX1 biological substrate. However, paradoxically, the TREX1 dominant mutant kinetics suggest that protomers in the TREX1 homodimer cannot simultaneously degrade large dsDNA substrates. To reconcile these findings, we propose a mechanism where TREX1 slides along the dsDNA backbone of its biological substrate in search of free 3’-hydroxyls, with each flexible loop “scanning” a respective DNA strand. Upon recognition and binding of a 3’-terminal nucleotide in the catalytic site of one TREX1 protomer by the α6 and/or α3-α4 regions, the flexible loop in the opposing protomer releases the non-scissile strand. The scissile DNA strand can then be positioned for efficient catalysis in the resident TREX1 protomer active site. This model helps to explain the ultra-fast TREX1 exonuclease kinetics (Tables 1–2), the disproportionate contribution of the TREX1 flexible loop to ds- versus ssDNA exonuclease activity (13), and the dominant TREX1 mutant kinetics (Figure 6). Further mechanistic studies are warranted to validate this model. Overall, our findings support a model where the TREX1 homodimer has evolved to efficiently locate free 3’-hydroxyls on a large, nicked dsDNA substrate and degrade short nucleotide stretches from the nicked DNA strand.

## Materials and methods

### Overexpression and purification of TREX1 enzymes

We have published a detailed protocol for purifying TREX1 enzymes (19), but we will summarize it here. Our experiments used truncated forms (aa. 1-242) of the human (UniProt: QN9SU2-3) and murine (UniProt: Q91XB0) TREX1 enzymes which contain only the catalytic domain, referred to in this work as ‘hT1’ and ‘mT1’, respectively. Wild-type protomers were expressed as a fusion protein using a pLM303x vector that encodes maltose-binding protein (MBP) linked N-terminally to TREX1. Mutant protomers were expressed as a fusion protein using a pCDF-Duet vector that encodes His-tagged NusA linked N-terminally to TREX1. Both constructs employed a linker with a rhinovirus 3C protease (PreScission Protease) recognition site and were transformed into *Escherichia coli* Rosetta2(DE3) cells (Novagen) for overexpression. Homodimers were purified via amylose column chromatography followed by overnight protease cleavage, then phosphocellulose (p-cell) column chromatography. Heterodimers were purified via amylose column chromatography followed by nickel column chromatography, then overnight protease cleavage followed by p-cell column chromatography. Amylose and nickel columns used one-step elution, and p-cell columns used a salt gradient for elution. All enzymes eluted from p-cell columns at a salt concentration of ~120 mM. Enzyme concentrations of preps were determined by A_280_ on a NanoDrop 2000 (Thermo Fisher) spectrophotometer and validated by SDS-PAGE.

### Approximation of TREX1 ssDNA k_cat_ and K_m_

Beginning from classic equations for the initial velocity of uninhibited enzyme (33), we derive Eq. 1 where *v_0_*, *k_cat_*, *[E_T_]*, *K_m_*, and *[S]* are initial velocity, catalytic rate constant, enzyme concentration, Michaelis-Menten constant, and substrate concentration, respectively.

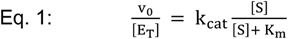

Deriving from classic (33) equations we obtain Eq. 2–3, where *k_1_* represents the association rate of enzyme and substrate (Eq. 4.1), *k_-1_* represents the dissociation rate of enzyme-substrate complex (Eq. 4.2), and *k_cat_* represents the catalytic rate constant for conversion of enzyme-substrate to enzyme and product (Eq. 4.3).

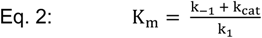

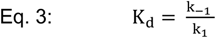

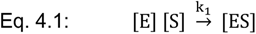

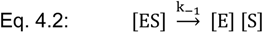

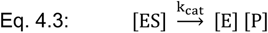

Beginning from Eq. 1, we rearrange to solve for Eq. 5 and 6.

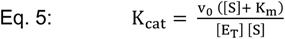

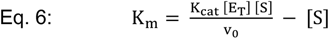

If the catalytic rate, *v_o_ / [E_T_]*, of the enzyme is determined at two substrate concentrations, then two iterations of Eq. 5 and 6 are obtained. Setting these iterations equal and solving for *K_m_* and *k_cat_* gives Eq. 7 and 8, respectively. We use *[S]_1_* and *[S]_2_* to differentiate the two substrate concentrations, and *β_1_* and *β_2_* to indicate the catalytic rates (*v_o_ / [E_T_]*) of the enzyme at those respective substrate concentrations.

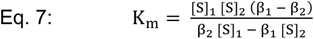

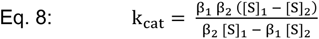

For this study ssDNA constants in Table 1, reported mean and standard deviation were approximated via Monte-Carlo simulation in R v3.6.1 (34) using a custom script. We generated vectors of 10^6^ values each for the *β_1_* and *β_2_* variables. Values were generated by uncorrelated sampling from a gamma distribution parameterized by shape = mean^2^/SD^2^ and rate = mean/SD^2^ using the wild-type values provided in Figure 2D. We used the substrate concentrations outlined in the ‘Polyacrylamide Gel ssDNA Assay’ Methods to define the *[S]_1_* and *[S]_2_* parameters. These vectors and parameters were applied pairwise to Eq. 7–8 to generate equal length vectors for *k_cat_* and *K_m_*. The *k_cat_* vector was then subjected to pairwise division by the *K_m_* vector to generate an equal length vector for *k_cat_*/*K_m_*. Means and standard deviations of the *k_cat_*, *K_m_*, and *k_cat_*/*K_m_* vectors were then calculated. For the *k_cat_*/*K_m_* values of previous studies, the *k_cat_* and *K_m_* vectors used to calculate the *k_cat_*/*K_m_* vector were generated by uncorrelated sampling from a gamma distribution as described above using the previously reported means and standard deviations (4).

### Absorbance-based dsDNA exonuclease assay

To generate a standard curve, synthetic dAMP nucleotides were serially diluted from 10 mM to 5 µM in 2-fold increments, then all dilution absorbances at 260 nm measured on a NanoDrop 2000 (Thermo Fisher) spectrophotometer. The plot of A_260_ vs [dAMP] was fit with linear regression in Prism 7.0 (GraphPad). The linear fit equation was rearranged and adjusted to the reported (35) extinction coefficient for dAMP (ε_260_ = 14,690 M^−1^cm^−1^) to produce Eq. 9. In this equation, *A_260_* indicates the absorbance at 260 nm, and *ε_dNMP_* indicates the average extinction coefficient of the nucleotides. The extinction coefficient parameter was approximated as the average of all reported (35) DNA nucleotide extinction coefficients, ε_260_ = 1160 M^−1^cm^−1^.

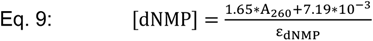

For the activity assays, 50 µL reactions were prepared containing 30 nM hT1, 20 mM TRIS (pH 7.5 @ 25°C), 5 mM MgCl_2_, 2 mM DTT, a variable concentration of dsDNA substrate, and 100 ng/µL BSA. The dsDNA substrate was a ~10-kb Nt.BbvCI-nicked (NEB #R0632) pMYC plasmid used at 2.5, 5, 10, 20, 40, and 75 nM. Reactions were initiated by addition of 10X enzyme diluted in 1 mg/mL BSA and incubated for 10 min at room temperature (~21°C). Reactions were quenched by the addition of 2.5 µL of 0.5 M EDTA. Quenched reactions were filtered with 10-kDa molecular weight cutoff microcentrifugal filter units according to manufacturer specifications (Amicon Ultra #UFC5010BK), then each flow-through had its absorbance measured at 260 nm on a NanoDrop 2000 (Thermo Fisher) spectrophotometer. Absorbance for each reaction was converted into concentration of excised nucleotides via Eq. 9, then used to calculate activity rate as moles of nucleotides excised per second. Plots of activity rate versus substrate concentration were fit with nonlinear regression in Prism 7.0 (GraphPad) consistent with Eq. 1 to solve for *k_cat_ [E_T_]* and *K_m_*.

### Polyacrylamide gel ssDNA assay

We have published a quite detailed protocol (19) for this assay, but we will summarize it here. 100 µL reactions were prepared containing a variable concentration of hT1, 20 mM TRIS (pH 7.5 @ 25°C), 5 mM MgCl_2_, 2 mM DTT, 15 nM or 515 nM ssDNA substrate, and 100 ng/µL BSA. Reactions were initiated by addition of 10X enzyme diluted in 1 mg/mL BSA and incubated for 20 minutes at room temperature (~21°C). Reactions were quenched in 400 µL cold ethanol, dried *in vacuo*, resuspended in loading solution, then electrophoresed on a 23% urea-polyacrylamide sequencing gel and imaged on a Typhoon FLA 9500 (GE Healthcare Life Sciences).

For the initial experiment to approximate TREX1 ssDNA k_cat_ and K_m_, the substrate was a 5’-FAM-labeled non-annealing 30-mer oligo of the below sequence, used at the indicated concentrations with the indicated concentrations of hT1. For the experiments to optimize the original kinetic model we proposed, the substrate was the same provided below used at 15 nM, and the enzyme was 10 pM of hT1. The optimization experiment included 6 independent reactions. In addition, a single reaction with an increased concentration of wild-type enzyme was used to generate a full ladder of oligo sizes for reference during quantification.

5’-FAM-labeled 30-mer oligo sequence: 5’-[FAM] ATACGACGGT GACAGTGTTG TCAGACAGGT-3’

### Quantification of polyacrylamide gels

Polyacrylamide gel images were subjected to 1D densitometry using the GelAnalyzer 2010a software. Lanes were identified automatically with software defaults, and intensity vs migration functions for each lane were subjected to background subtraction by the rolling ball method with a ‘500’ ball radius setting. Next, the peaks in each lane corresponding to various oligo sizes were manually identified using the aforementioned internal ladder as a visual reference for the expected migration positions. Then, peaks were integrated automatically to quantify relative oligo levels for all bands. These band values were subjected to quantification methods (19) we have previously published to calculate the proportion of each substrate size remaining in the reaction, calculate the moles of nucleotides excised, and ultimately determine reaction rates.

### Thermal shift assays

For the assay, 100 µL reactions were prepared containing 0.5 mg/mL TREX1, 20 mM TRIS (pH 7.5 @ 25°C), 5 mM CaCl_2_, 2 mM DTT, 5X SYPRO Orange dye (ThermoFisher #S6650), and a variable concentration of substrate. No BSA was included. The dsDNA substrate was a self-annealing 30-mer oligonucleotide, and the ssDNA substrate was a 30-mer oligonucleotide. Both substrates were included at 18 µM in the relevant reactions. 8 replicate reactions per condition/enzyme were created, reconciled into a clear 96-well PCR plate (VWR #76402-848), and sealed with adhesive plate tape. Melting curve experiments were then performed with a 7500 Real-Time PCR System (Applied Biosystems). Melting experiment was conducted over a 25-95°C temperature range, at a rate of 1°C/min. Fluorescence was monitored with the system TAMRA reporter settings, with 188 fluorescence readings per sample in equal ΔT intervals across the temperature gradient.

Melting curve data was imported to R, then each sample curve normalized internally to maximum and minimum fluorescence. Curves were then fit with a smoothing spline using the smooth.spline function (stats package). Fit curves were then used to interpolate T_m_ from the melting curve inflections points, via the deriv function (stats package). All calculations described in this section were performed with R v3.6.1 (34).

30-mer oligo sequence (dsDNA): 5’-GCTCGAGTCA TGACGCGTCA TGACTCGAGC-3’

30-mer oligo sequence (ssDNA): 5’-TTAACCTTCT TTATAGCCTT TGAACAAAGG-3’

### Molecular dynamics (MD) simulations

The published structure of mTREX1_1-242_ bound to ssDNA (PDB = ‘2IOC’) was modified in PyMOL (36) to remove nucleic acid and ions. The modified structure was then used as a template for MODELLER (37) to model in the flexible loops missing from the original structure. This produced the mT1 model. The original mTREX1 published structure was loaded into PyMOL (36) and aligned to the mT1 model. The calcium ions and ssDNA polymers from the original structure were then superimposed on the mT1 model, and the calcium ions converted to magnesium, to generate the mT1:4-mer model. The published structure of mTREX1_1-242_ bound to dsDNA (PDB = ‘4YNQ’) was loaded into PyMOL (36) and aligned to the mT1:4-mer model. The ssDNA polymers were removed from the mT1:4-mer model, a single dsDNA from the published structure copied into the model, the published structure removed, and then the remaining molecules exported. This generated the mT1:dsDNA model.

Each of the models was converted to pdb and psf files with the autopsf functionality in VMD (v1.9.4) (38), using the CHARMM top_all27_prot_lipid_na.inp force field parameters (39, 40). Explicit water solvent was added to the processed models using a 10 Å padding around the existing model dimensions. Hydrated models were then ionized with 50 mM MgCl_2_ and exported. The exported pdb and psf files defined the initial conditions for the four simulation systems, constructed from the four models. Hydration and ionization were also performed in VMD v1.9.4 using default CHARMM parameter files.

Initial condition files for each system were replicated and used to initiate four MD simulations per system. MD simulations were performed with ACEMD3 (Acellera) (41) on Acellera GPUs. All initial condition files were independently subjected to a 1000-step energy minimization, then simulations were initiated. Simulations were for 1000 ns each, with 4 fs steps. Structure coordinates were exported from simulations every 10 ps in xtc format. Thermostat was set to 300 K, barostat was set to 1.0132 bar, and all other parameters were left to default. Simulations ran for ~10 days each.

Completed simulations were pre-processed in VMD v1.9.4. The xtc trajectory file and psf initial conditions file were loaded into the pdb initial conditions file. Simulation trajectory coordinates were loaded in 20-frame (200 ps) intervals totaling 5000 frames. After loading, simulations were unwrapped using the pbc unwrap function. Unwrapped simulation frames were aligned to the initial conditions frame based on RMSD-minimization between the protein backbones. Pre-processed simulations were exported without solvent molecules to mol2 files. The mol2 files were processed into csv files compatible with later analyses using Bash scripts.

### Calculating phi and psi angles of TREX1 residues

The python package MDAnalysis was used to calculate phi and psi angles for the TREX1 residues. For each simulation of each system, the respective pre-processed mol2 file was analyzed with the Dihedral and Ramachandran functions to generate phi and psi angles for every time-point (frame) of the simulations. Phi and psi values were then exported as separate csv files compatible with subsequent analyses. Python script for phi/psi angle calculations was run with python v3.7.4.

### Calculating RMSD

The RMSD values for each simulation were calculated via Eq. 10. Where: ***r****_j_(t_i_)* is the position vector of the j^th^ atom in the i^th^ time frame and N is the total number of atoms.

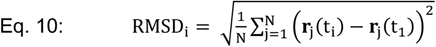

These calculations provided RMSD values at every time frame for all four simulations per system. The values for all simulations were averaged by time frame to give mean values. All calculations described in this section were performed in R v3.6.1 (34).

### Calculating RMSF

The RMSF values in each simulation were calculated via Eq. 11. Where: where ***r****_i_(t_j_)* is the instantaneous position vector of the i^th^ atom, ***r’****_i_* is the mean position of the i^th^ atom, and T is the total number of time frames.

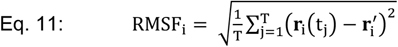

The RMSF values for all simulations of a system were grouped by atom to give RMSF means and standard deviations. All calculations described in this section were performed in R v3.6.1 (34).

### Clustering by amino acid phi and psi angles

The phi and psi values were first converted into Cartesian coordinates via projection onto a sphere with Eq. 12–14. This allowed seamless clustering for residue conformations that had phi (Ψ) or psi (Φ) values spanning the −180°/180° transition.

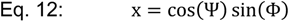

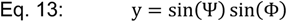

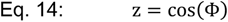

For each residue, every third time frame of every simulation of every system was clustered by these projected coordinates using the hdbscan function (dbscan package) (42) in R with a minPts=200 parameter to attribute any clusters containing <1% of time frames to noise. For every residue, the proportion of each system time frames belonging to the identified clusters/conformations were calculated and pairwise comparisons of the proportions between systems were made via a chi-squared test of independence, using the chisq.test function (stats package) in R. P-values from these comparisons were converted to q-values by adjusting for false discovery rate with the Benjamini-Hochberg (43) method via the p.adjust function (stats package) in R. Residues with significantly different conformation proportions between systems were then identified. All calculations and statistics for this section were performed in R v3.6.1 (34).

### Conformational free energy maps

The coordinates of every relevant atom were used as variables in a principal components analysis (PCA) on every time frame of every simulation of every relevant system. The PCA was performed via the ‘princomp()’ function in R. Each resulting principal component (PC) proportion of variance (PV) was calculated via Eq. 15, where *σ_i_^2^* is the variance for the i^th^ principal component and *N* is the number of principal components.

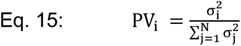

Values for the first two PCs were extracted and separated by system. Density functions for each system were then approximated using PC-1 and PC-2 with the kde2d function (MASS package) (44) in R. Density functions for each system were visualized as contour plots using the contour function (graphics package) in R. All calculations and statistics for this section were performed in R v3.6.1 (34).

### Kinetic modeling

For Eq. 16.1–5, apply Figure 3 notation, *S_β:n_* is all substrates with between *β* and *n* excisable nucleotides, *β* is the maximum number of excisable nucleotides on a substrate molecule during the reaction, 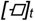 is concentration of the respective reactant or complex at time *t*, 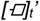 is rate of change in concentration of the respective reactant or complex at time *t*, and all other previous notation applies. Reference to Figure 3 scheme 1 provides helpful context.

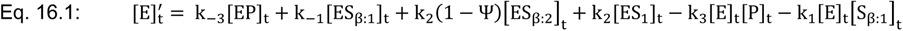

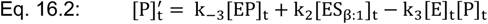

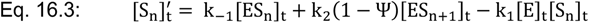

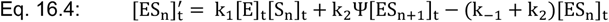

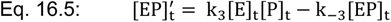

For Eq. 17.1–8, apply Figure 3 notation, *[E]* and *[M]* are concentrations of free wild-type and mutant enzyme, respectively, *Ψ* and *Φ* are the wild-type and mutant processivity parameters, respectively, and all other notation is previously established. Reference to Figure 3 scheme 2 provides useful context.

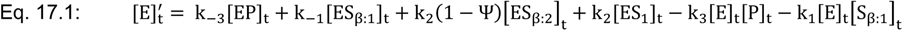

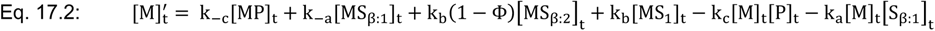

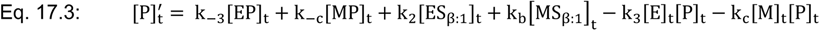

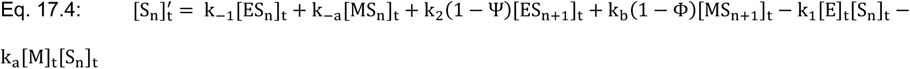

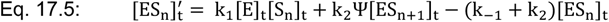

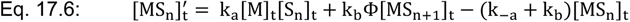

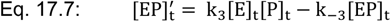

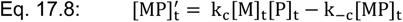

For Eq. 18, apply Figure 3 notation, and *P_Ψ_*(*N*) is the probability of *N* nucleotides being excised before dissociation with a processivity factor of *Ψ*.

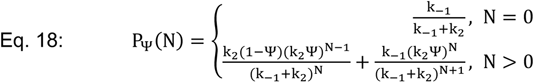

*Assuming 0^0^ = 1, as opposed to being undefined.

For Eq. 19, apply classic (33) notation.

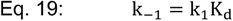

In both schemes (Figure 3), turnover rate (Eq. 16.2) reduces to k_2_ [E_T_] under saturating substrate conditions if enzyme-product complexing is negligible, so the TREX1 catalytic rate constants (k_2_) for dsDNA and ssDNA are equivalent to k_cat_ (33) from our steady-state kinetics studies (Table 1). The previously reported (21) ssDNA K_d_ value for TREX1 is applicable to our model with the classic (33) relationship of Eq. 19. Using the constraint of Eq. 19, we compared our ssDNA exonuclease activity rates (Figure 2D, [S] = 15 nM) with numerical solutions to Eq. 16.1–5 to approximate the on-rate constant (k_1_) and off-rate constant (k_-1_) values required to recapitulate our data for various processivity parameter (Ψ) values (Supplemental 1). We then compared our empirical ssDNA degradation patterns (Figure 2D, [S] = 15 nM) to those predicted by these various rate constant solution sets (Figure 4A).

For dsDNA, data from hT1 and hT1-DN competition experiments were compared with numerical solutions to Eq. 17.1–8 for a range of fixed processivity parameter values to determine the corresponding solutions for the TREX1 dsDNA on-rate and off-rate constants (Supplemental 2). To capture hT1-DN biochemistry in our modeling, the hT1-DN catalytic constant was set to zero and all other kinetic constants were constrained to wild-type values. Most processivity constant values had solutions for k_1_ and k_-1_ that could recapitulate our data, but for Ψ ≥ 0.90 there were no solutions that well recapitulated our data (Supplemental 2D-F). Under the additional assumption that TREX1 exhibits equal or greater processivity on ds- versus ssDNA (Ψ_ds_ ≥ Ψ_ss_), we can apply the TREX1 dsDNA rate constant constraints of 0.75 ≤ Ψ ≤ 0.90, 0.020 ≤ k_-1_ ≤ 0.10 s^−1^, and k_1_ ≈ 10^9^ M^−1^s^−1^ (Table 2). Rate constant solutions for Ψ = 0 were similar (k_1_ = 5.0×10^9^ M^−1^s^−1^, k_-1_ = 0.30 s^−1^).

We provide detailed accounts of the parameter definitions for every simulated reaction (Supplemental 6), which should allow reproduction of these simulations. Any parameters not provided in the tables were set to ‘0’ under all conditions. Every reaction proposed by the kinetic model was simulated with the R script provided on GitHub.

### Fluorescence-based dsDNA exonuclease assay

We have published a detailed protocol (19) for this assay, but we will summarize it here. 150 µL reactions were prepared containing a variable concentration of hT1, 20 mM TRIS (pH 7.5 @ 25°C), 5 mM MgCl_2_, 2 mM DTT, 5-10 ng/µL dsDNA substrate, and 200 ng/µL BSA. For the ‘[S]<K_M_’ experiments, the dsDNA substrate was a ~10-kb Nt.BbvCI-nicked (NEB #R0632) pMYC plasmid used at 5 ng/µL (~0.83 nM), and the hT1 concentration was 15 nM. For the ‘[S]>>K_M_’ experiments, the dsDNA substrate was a self-annealing 30-mer oligo used at 1000 nM (~9 ng/µL), and the hT1 concentration was 2.5 nM. Reactions were initiated by addition of 10X enzyme diluted in 1 mg/mL BSA and incubated for 1 hour at room temperature (~21°C). 20 µL samples were taken at 0, 5, 10, 20, 30, 45, and 60-minute time points and quenched in 20 µL of 15X SYBR Green dye (ThermoFisher #S7563). Quenched samples had their fluorescence measured on a PolarStar Omega microplate reader (BMG LabTech), using excitation/emission of 485/520 nm. For the 1:1:0 enzyme mixes, mix ratios indicate the prevalence of hT1^+/+^, hT1^MUT/MUT^, and hT1^MUT/+^, respectively, where total hT1 concentrations were still as indicated.

30-mer oligo sequence: 5’-GCTCGAGTCA TGACGCGTCA TGACTCGAGC-3’

### Quantification of fluorescence-based data

Each independent enzyme dilution in each experiment was used to create 6 replicate reactions. Plots of fluorescence vs time were generated from the time-course data for every reaction in each experiment. The data points from every reaction replicate were used for fitting with one-phase decay nonlinear regression. This gave 4 fitted datasets per experiment, one for each independent enzyme dilution. The slope of each fit at 0 min was interpolated to give initial velocities, and then initial velocities were normalized within each experiment to the mean of the wild-type fit initial velocities to calculate relative activities for each variant/mix. Calculations were performed in Excel, and regression and interpolation were performed in Prism 9.0 (GraphPad).

### Cα correlation analyses

The Cα coordinates across all time frames and simulations were reconciled for each system, and Eq. 20.1–2 were used to calculate the correlation factors between atom pairs. Here, *C_ij_* is the correlation factor for atoms *i* and *j*, *N* is the number of frames, 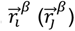 is the position of atom *i* (*j*) in time frame *β*, and 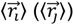 is the mean position or centroid of atom *i* (*j*) across all time frames. All calculations described in this section were performed in R v3.6.1 (34).

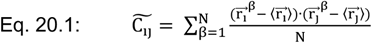

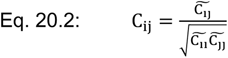

### Availability of data and scripts used for computational analyses

Scripts used for analyses described in these methods are available on GitHub (https://github.com/whemphil/TREX1_Modeling_Scripts.git), pre-processed data input files for MD analysis scripts have been deposited to Zenodo (doi.org/10.5281/zenodo.6347909), and raw files from MD simulations are available from W.O.H. upon request (total data ≈ 0.5 TB). Any other raw data requests should be directed to W.O.H.

## Supporting Information

This article contains supporting information.

## Acknowledgements

We’d like to thank the WFU high-performance computing team for access to the DEAC cluster and technical support.

## Author Contributions

W.O.H.: Conceptualization, Investigation, Methodology, Software, Visualization, Validation, Writing – Original draft preparation. T.H.: Writing – Reviewing and editing. F.R.S.: Conceptualization, Methodology, Software, Supervision, Writing – Reviewing and editing. F.W.P.: Conceptualization, Methodology, Supervision, Writing – Reviewing and editing. All authors approved the final manuscript.

## Funding & Additional Information

This work was supported by the National Institute of Allergy and Infectious Diseases (AI116725 to F.W.P.), National Institute of General Medical Sciences (GM110734 to F.W.P.), Lupus Research Alliance (PI – F.W.P.), Wake Forest Innovations (PI – F.W.P.), Comprehensive Cancer Center of Wake Forest University National Cancer Institute (Center Support Grant, P30CA012197), NIH NRSA Predoctoral Fellowships (T32-GM095440 & T32-AI007401; Trainee – W.O.H.), a Sandy Lee Cowgill Memorial Scholarship (Trainee – W.O.H.), an Artom Fellowship (Trainee – W.O.H.), and a Scott Family Fellowship (PI – F.R.S.). W.O.H. has been partially supported by the Howard Hughes Medical Institute (PI – Thomas Cech) and National Institutes of Health (F32-GM147934; PI – W.O.H.) during manuscript writing. The content is solely the responsibility of the authors and does not necessarily represent the official views of the National Institutes of Health.

## Abbreviations

AGS: Aicardi-Goutieres syndrome
cGAS: cyclic GMP-AMP synthase
Cα: alpha carbon
dNMP: deoxynucleoside monophosphate
FCL: familial Chilblain lupus
hT1: recombinant human TREX1_1-242_ enzyme
hT1-DN: recombinant humanTREX1_1-242_^D18N/D18N^ enzyme
hT1-DNhet: recombinant human TREX1_1-242_^WT/D18N^ enzyme
MD: molecular dynamics
mT1: recombinant murine TREX1_1-242_ enzyme
PC: principal component
PCA: principal components analysis
PV: proportion of variance
p-cell: phosphocellulose
RMSD: root mean squared deviation
RMSF: root mean squared fluctuation
RVCL: retinal vasculopathy with cerebral leukodystrophy
SLE: systemic lupus erythematosus
STING: stimulator of interferon genes
TREX1: three-prime repair exonuclease 1
TREX2: three-prime repair exonuclease 2

## Accession Codes

mTREX1: Q91XB0 (UniProt)

hTREX1: QN9SU2-3 (UniProt)

## References

1. Simpson, S. R., Hemphill, W. O., Hudson, T., and Perrino, F. W. (2020) TREX1 – Apex predator of cytosolic DNA metabolism. DNA Repair. 10.1016/j.dnarep.2020.102894

2. Rice, G. I., Rodero, M. P., and Crow, Y. J. (2015) Human Disease Phenotypes Associated With Mutations in TREX1. J. Clin. Immunol. 35, 235–243

3. Perrino, F. W., Miller, H., and Ealey, K. A. (1994) Identification of a 3’-->5’-exonuclease that removes cytosine arabinoside monophosphate from 3’ termini of DNA. J. Biol. Chem. 269, 16357–16363

4. Mazur, D. J., and Perrino, F. W. (2001) Excision of 3′ Termini by the Trex1 and TREX2 3′→5′ Exonucleases CHARACTERIZATION OF THE RECOMBINANT PROTEINS. J. Biol. Chem. 276, 17022–17029

5. Belyakova, N. V., Kleiner, N. E., Kravetskaya, T. P., Legina, O. K., Naryzhny, S. N., Perrino, F. W., Shevelev, I. V., and Krutyakov, V. M. (1993) Proof-reading 3’-->5’ exonucleases isolated from rat liver nuclei. Eur. J. Biochem. 217, 493–500

6. Mazur, D. J., and Perrino, F. W. (1999) Identification and Expression of the TREX1 and TREX2 cDNA Sequences Encoding Mammalian 3′→5′ Exonucleases. J. Biol. Chem. 274, 19655–19660

7. Ablasser, A., Hemmerling, I., Schmid-Burgk, J. L., Behrendt, R., Roers, A., and Hornung, V. (2014) TREX1 Deficiency Triggers Cell-Autonomous Immunity in a cGAS-Dependent Manner. J. Immunol. 192, 5993–5997

8. Gray, E. E., Treuting, P. M., Woodward, J. J., and Stetson, D. B. (2015) cGAS is required for lethal autoimmune disease in the Trex1-deficient mouse model of Aicardi-Goutieres Syndrome. J. Immunol. Baltim. Md 1950. 195, 1939–1943

9. Xiao, N., Wei, J., Xu, S., Du, H., Huang, M., Zhang, S., Ye, W., Sun, L., and Chen, Q. (2019) cGAS activation causes lupus-like autoimmune disorders in a TREX1 mutant mouse model. J. Autoimmun. 100, 84–94

10. Wu, J., Sun, L., Chen, X., Du, F., Shi, H., Chen, C., and Chen, Z. J. (2013) Cyclic-GMP-AMP Is An Endogenous Second Messenger in Innate Immune Signaling by Cytosolic DNA. Science. 10.1126/science.1229963

11. Rice, G., Newman, W. G., Dean, J., Patrick, T., Parmar, R., Flintoff, K., Robins, P., Harvey, S., Hollis, T., O’Hara, A., Herrick, A. L., Bowden, A. P., Perrino, F. W., Lindahl, T., Barnes, D. E., and Crow, Y. J. (2007) Heterozygous Mutations in TREX1 Cause Familial Chilblain Lupus and Dominant Aicardi-Goutières Syndrome. Am. J. Hum. Genet. 80, 811–815

12. Bailey, S. L., Harvey, S., Perrino, F. W., and Hollis, T. (2012) Defects in DNA degradation revealed in crystal structures of TREX1 exonuclease mutations linked to autoimmune disease. DNA Repair. 11, 65–73

13. Fye, J. M., Orebaugh, C. D., Coffin, S. R., Hollis, T., and Perrino, F. W. (2011) Dominant Mutations of the TREX1 Exonuclease Gene in Lupus and Aicardi-Goutières Syndrome. J. Biol. Chem. 286, 32373–32382

14. Orebaugh, C. D., Fye, J. M., Harvey, S., Hollis, T., Wilkinson, J. C., and Perrino, F. W. (2013) The TREX1 C-terminal Region Controls Cellular Localization through Ubiquitination. J. Biol. Chem. 288, 28881–28892

15. Orebaugh, C. D., Fye, J. M., Harvey, S., Hollis, T., and Perrino, F. W. (2011) The TREX1 Exonuclease R114H Mutation in Aicardi-Goutières Syndrome and Lupus Reveals Dimeric Structure Requirements for DNA Degradation Activity♦. J. Biol. Chem. 286, 40246–40254

16. Silva, U. de, Choudhury, S., Bailey, S. L., Harvey, S., Perrino, F. W., and Hollis, T. (2007) The Crystal Structure of TREX1 Explains the 3′ Nucleotide Specificity and Reveals a Polyproline II Helix for Protein Partnering. J. Biol. Chem. 282, 10537–10543

17. Fye, J. M., Coffin, S. R., Orebaugh, C. D., Hollis, T., and Perrino, F. W. (2014) The Arg-62 residues of the TREX1 exonuclease act across the dimer interface contributing to catalysis in the opposing protomers. J. Biol. Chem. 289, 11556–11565

18. Hall, J. (2019) A simple model for determining affinity from irreversible thermal shifts. Protein Sci. Publ. Protein Soc. 28, 1880–1887

19. Hemphill, W. O., and Perrino, F. W. (2019) Measuring TREX1 and TREX2 exonuclease activities. Methods Enzymol. 625, 109–133

20. Lindahl, T., Gally, J. A., and Edelman, G. M. (1969) Properties of Deoxyribonuclease III from Mammalian Tissues. J. Biol. Chem. 244, 5014–5019

21. de Silva, U. (2007) *STRUCTURAL AND BIOCHEMICAL STUDIES OF TREX - THREE PRIME REPAIR EXONUCLEASES*. Ph.D. thesis, WAKE FOREST UNIVERSITY GRADUATE SCHOOL OF ARTS AND SCIENCES

22. Bailey, S. L. (2010) The TREX1 3’exonuclease and autoimmune disease: Structural and biochemical analysis of disease mutants involved in autoimmune dysfunction. Ph.D. thesis, Wake Forest University

23. Grieves, J. L., Fye, J. M., Harvey, S., Grayson, J. M., Hollis, T., and Perrino, F. W. (2015) Exonuclease TREX1 degrades double-stranded DNA to prevent spontaneous lupus-like inflammatory disease. Proc. Natl. Acad. Sci. U. S. A. 112, 5117–5122

24. Perrino, F. W., de Silva, U., Harvey, S., Pryor, E. E., Cole, D. W., and Hollis, T. (2008) Cooperative DNA Binding and Communication across the Dimer Interface in the TREX2 3′ → 5′-Exonuclease. J. Biol. Chem. 283, 21441–21452

25. Lehtinen, D. A., Harvey, S., Mulcahy, M. J., Hollis, T., and Perrino, F. W. (2008) The TREX1 Double-stranded DNA Degradation Activity Is Defective in Dominant Mutations Associated with Autoimmune Disease. J. Biol. Chem. 283, 31649–31656

26. Fersht, A. (1985) Enzyme Structure and Mechanism, 2nd Ed., W. H. Freeman and Co., New York

27. de Silva, U., Perrino, F. W., and Hollis, T. (2009) DNA binding induces active site conformational change in the human TREX2 3′-exonuclease. Nucleic Acids Res. 37, 2411–2417

28. Berg, O. G., Winter, R. B., and von Hippel, P. H. (1981) Diffusion-driven mechanisms of protein translocation on nucleic acids. 1. Models and theory. Biochemistry. 20, 6929–6948

29. Hemphill, W. O., Voong, C. K., Fenske, R., Goodrich, J. A., and Cech, T. R. (2022) RNA- and DNA-binding proteins generally exhibit direct transfer of polynucleotides: Implications for target site search. 10.1101/2022.11.30.518605

30. Fazlieva, R., Spittle, C. S., Morrissey, D., Hayashi, H., Yan, H., and Matsumoto, Y. (2009) Proofreading exonuclease activity of human DNA polymerase δ and its effects on lesion-bypass DNA synthesis. Nucleic Acids Res. 37, 2854–2866

31. Huang, K.-W., Hsu, K.-C., Chu, L.-Y., Yang, J.-M., Yuan, H. S., and Hsiao, Y.-Y. (2016) Identification of Inhibitors for the DEDDh Family of Exonucleases and a Unique Inhibition Mechanism by Crystal Structure Analysis of CRN-4 Bound with 2-Morpholin-4-ylethanesulfonate (MES). J. Med. Chem. 59, 8019–8029

32. Mason, P. A., and Cox, L. S. (2012) The role of DNA exonucleases in protecting genome stability and their impact on ageing. Age. 34, 1317–1340

33. Ault, A. (1974) An introduction to enzyme kinetics. J. Chem. Educ. 51, 381

34. R Core Team (2021) R: A language and environment for statistical computing.

35. Onidas, D., Markovitsi, D., Marguet, S., Sharonov, A., and Gustavsson, T. (2002) Fluorescence Properties of DNA Nucleosides and Nucleotides: A Refined Steady-State and Femtosecond Investigation. J. Phys. Chem. B. 106, 11367–11374

36. Schrodinger, L. and DeLano, W. (2020) PyMOL

37. Webb, B., and Sali, A. (2016) Comparative Protein Structure Modeling Using MODELLER. Curr. Protoc. Bioinforma. Ed. Board Andreas Baxevanis Al. 54, 5.6.1–5.6.37

38. Humphrey, W., Dalke, A., and Schulten, K. (1996) VMD: Visual molecular dynamics. J. Mol. Graph. 14, 33–38

39. Foloppe, N., and MacKerell, A. D., Jr. (2000) All-atom empirical force field for nucleic acids: I. Parameter optimization based on small molecule and condensed phase macromolecular target data. J. Comput. Chem. 21, 86–104

40. MacKerell, A. D., Bashford, D., Bellott, M., Dunbrack, R. L., Evanseck, J. D., Field, M. J., Fischer, S., Gao, J., Guo, H., Ha, S., Joseph-McCarthy, D., Kuchnir, L., Kuczera, K., Lau, F. T., Mattos, C., Michnick, S., Ngo, T., Nguyen, D. T., Prodhom, B., Reiher, W. E., Roux, B., Schlenkrich, M., Smith, J. C., Stote, R., Straub, J., Watanabe, M., Wiórkiewicz-Kuczera, J., Yin, D., and Karplus, M. (1998) All-atom empirical potential for molecular modeling and dynamics studies of proteins. J. Phys. Chem. B. 102, 3586–3616

41. Harvey, M. J., Giupponi, G., and Fabritiis, G. D. (2009) ACEMD: Accelerating Biomolecular Dynamics in the Microsecond Time Scale. J. Chem. Theory Comput. 5, 1632–1639

42. Hahsler, M., Piekenbrock, M., and Doran, D. (2019) dbscan: Fast Density-Based Clustering with R. J. Stat. Softw. 91, 1–30

43. Benjamini, Y., and Hochberg, Y. (1995) Controlling the False Discovery Rate: A Practical and Powerful Approach to Multiple Testing. J. R. Stat. Soc. Ser. B Methodol. 57, 289–300

44. Venables, W.N. and Ripley, B.D. (2002) Modern Applied Statistics with R, Fourth, Springer New York

