## Supporting Information for "A TREX1 model reveals double-strand DNA preference and inter-protomer regulation"

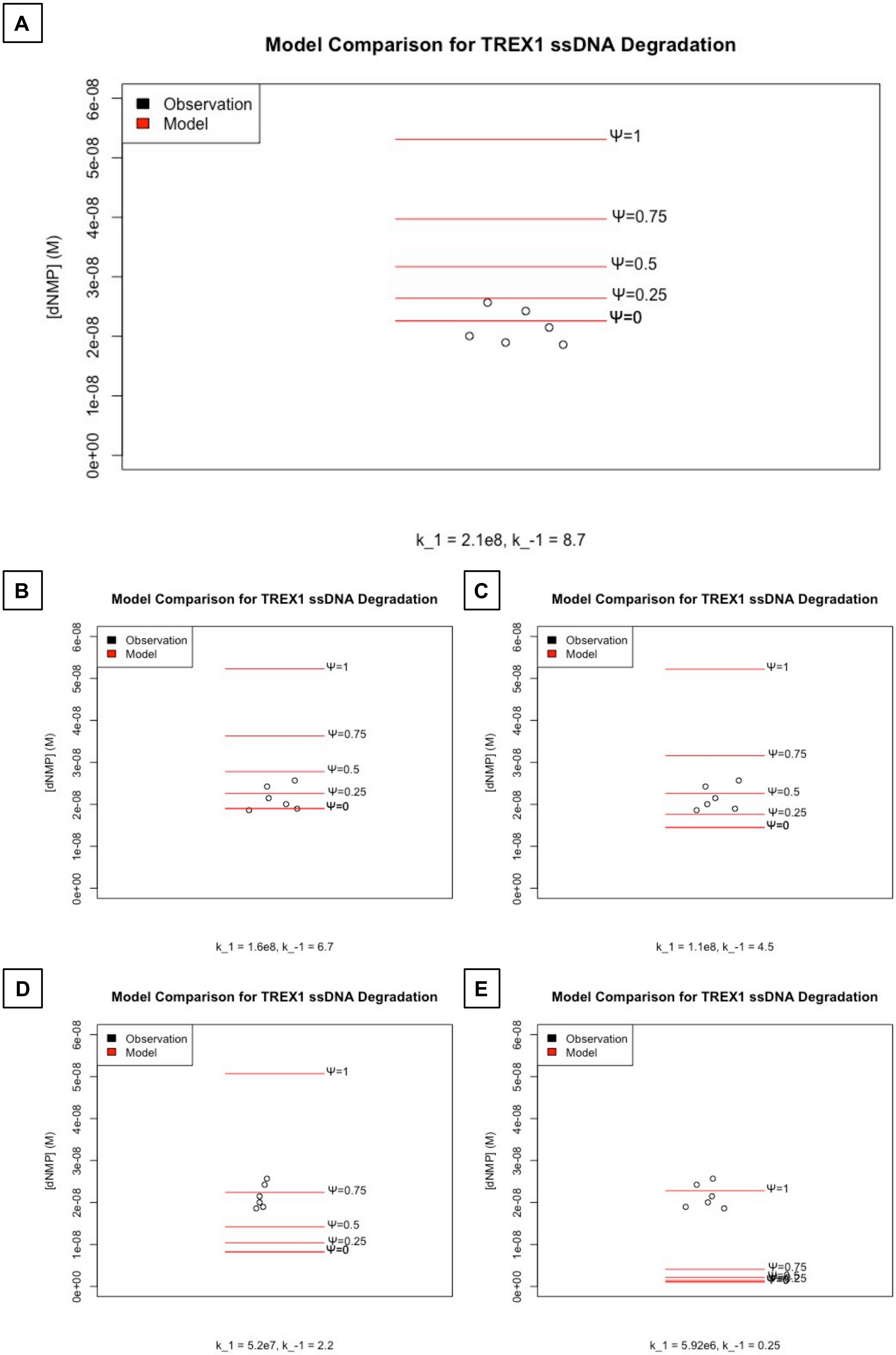

A
Model Comparison for TREX1 ssDNA Degradation
Observation
Model
[dNMP] (M)
0e+00
1e-08
2e-08
3e-08
4e-08
5e-08
6e-08
Ψ=1
Ψ=0.75
Ψ=0.5
Ψ=0.25
Ψ=0
k_1 = 2.1e8, k_-1 = 8.7
B
Model Comparison for TREX1 ssDNA Degradation
Observation
Model
[dNMP] (M)
Ψ=1
Ψ=0.75
Ψ=0.5
Ψ=0.25
Ψ=0
k_1 = 1.6e8, k_-1 = 6.7
C
Model Comparison for TREX1 ssDNA Degradation
Observation
Model
[dNMP] (M)
Ψ=1
Ψ=0.75
Ψ=0.5
Ψ=0.25
Ψ=0
k_1 = 1.1e8, k_-1 = 4.5
D
Model Comparison for TREX1 ssDNA Degradation
Observation
Model
[dNMP] (M)
Ψ=1
Ψ=0.75
Ψ=0.5
Ψ=0.25
Ψ=0
k_1 = 5.2e7, k_-1 = 2.2
E
Model Comparison for TREX1 ssDNA Degradation
Observation
Model
[dNMP] (M)
Ψ=1
Ψ=0.75
k_1 = 5.92e6, k_-1 = 0.25

**<u>Supplemental 1</u>**: **Optimization of parameters for TREX1 ssDNA kinetic model.** A polyacrylamide gel-based TREX1 ssDNA degradation assay was performed as described in the relevant methods section, and the banding data quantified to give the concentration of product in the reaction. Product concentrations for all 6 replicate reactions are displayed ('Observation'). Any variability along the x-axis in the empirical data points between graphs is not indicative of a different data set but an artifact of R plotting code. The kinetic model for hTREX1 ssDNA exonuclease activity was optimized at several processivity parameter values ('Ψ') to determine the matching kinetic constant values ('k_1', 'k_-1') that gave the best fit to the empirical data. The 'A-E' plots correspond to optimization for Ψ values of 0, 0.25, 0.5, 0.75, and 1, respectively. Labels on the bottom-x axis of each graph indicate the model relevant reaction parameters, and the labeled red lines on each graph indicate the predicted product concentration for indicated Ψ values under those parameters. All calculations and plotting were performed in R v3.6.1, and figure preparation was performed in PowerPoint (Microsoft). Simulated reactions are defined in Supplemental 6.

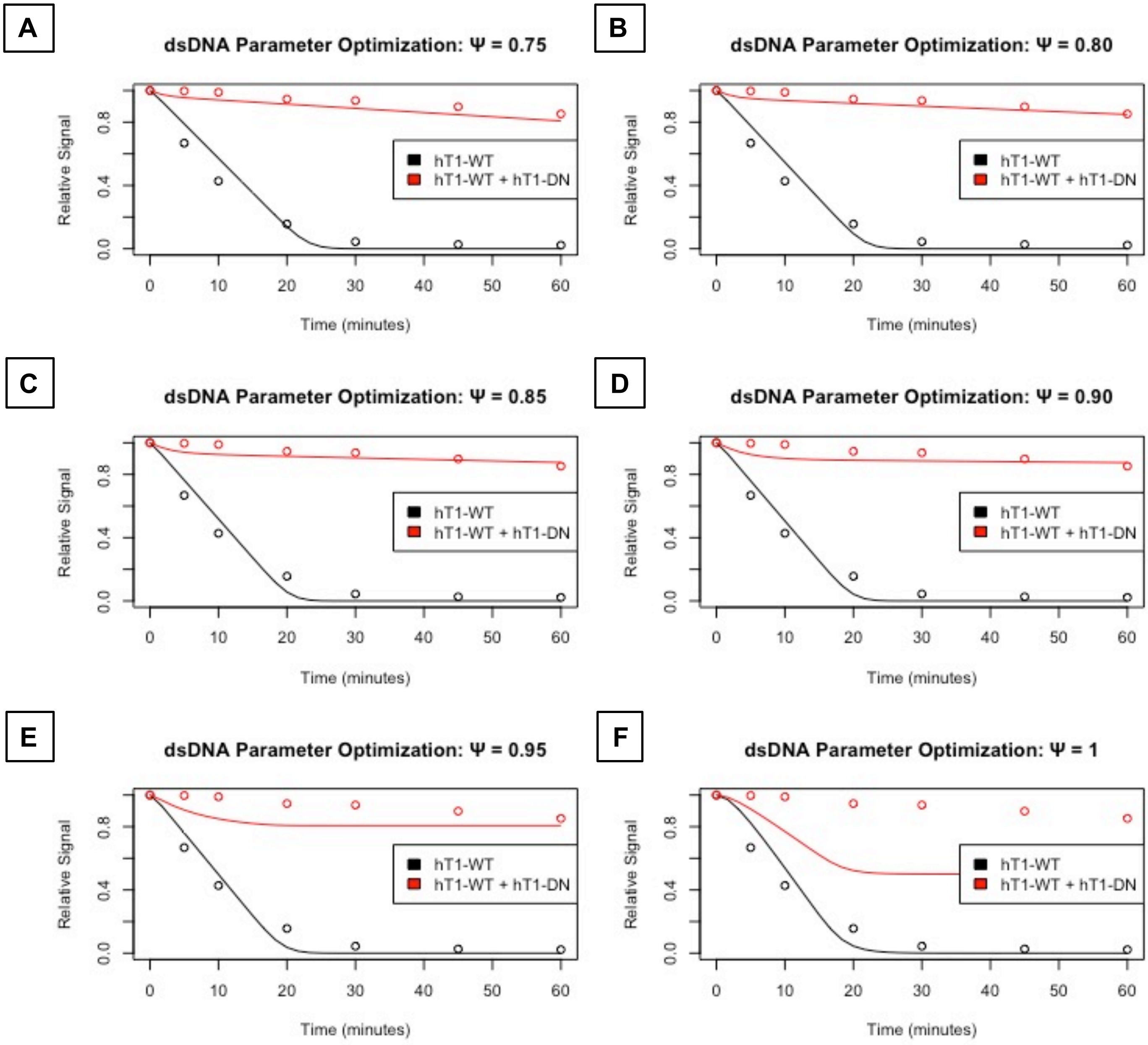

A
dsDNA Parameter Optimization: Ψ = 0.75
B
dsDNA Parameter Optimization: Ψ = 0.80
C
dsDNA Parameter Optimization: Ψ = 0.85
D
dsDNA Parameter Optimization: Ψ = 0.90
E
dsDNA Parameter Optimization: Ψ = 0.95
F
dsDNA Parameter Optimization: Ψ = 1
Relative Signal
Time (minutes)
0.0
0.4
0.8
0
10
20
30
40
50
60
hT1-WT
hT1-WT + hT1-DN

**<u>Supplemental 2</u>**: **Optimization of parameters for TREX1 dsDNA kinetic model.** Standard exonuclease reactions were prepared with 15 nM hT1 (black data) or 15 nM of a 1:1 mix of hT1:hT1-DN (red data), incubated at room temperature for the indicated times, quenched in SYBR Green, and dsDNA content measured by fluorescence. Plots of fluorescence vs time were generated and normalized to maximum and minimum fluorescence. Data points on the graph correspond to these empirical data, and they represent means of 18 reactions per condition across three experiments. The parameters of the TREX1 dsDNA kinetic model were optimized to fit these empirical data for each of the processivity values listed in respective plot titles. Optimization was carried out by starting with the ssDNA $k_1$ and $k_{-1}$ values, increasing $k_1$ until the predicted wild-type degradation rate (black lines) matched observation, then decreasing $k_{-1}$ until the predicted level of enzyme competition (red lines) matched observation. Calculations and plotting were performed in R v3.6.1. Simulated reactions are defined in Supplemental 6. The black and red data points here are derived from the same data as the blue and purple data points in Figure 6A, respectively, after normalization to maximum and minimum fluorescence.

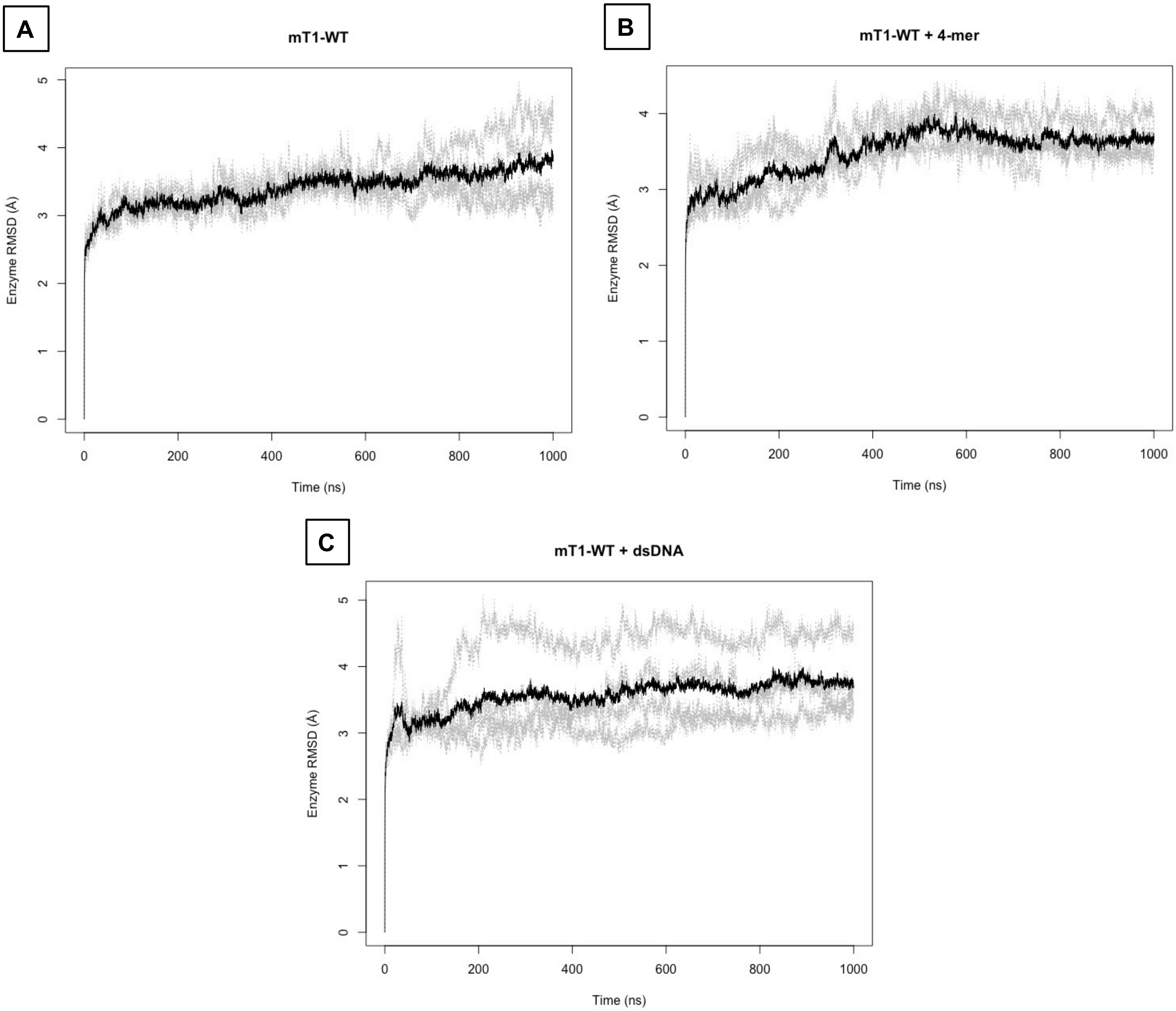
A
mT1-WT
Enzyme RMSD (Å)
Time (ns)
B
mT1-WT + 4-mer
Enzyme RMSD (Å)
Time (ns)
C
mT1-WT + dsDNA
Enzyme RMSD (Å)
Time (ns)

**<u>Supplemental 3</u>**: **RMSD of mTREX1 structure from initial conditions.** All TREX1 atoms were used in an RMSD calculation as described in the relevant methods section. RMSD from initial conditions was plotted as a function of time. Grey lines indicate individual simulations, and black lines indicate averages for the 4 replicate simulations of each system. All calculations and graphing were performed in R v3.6.1.

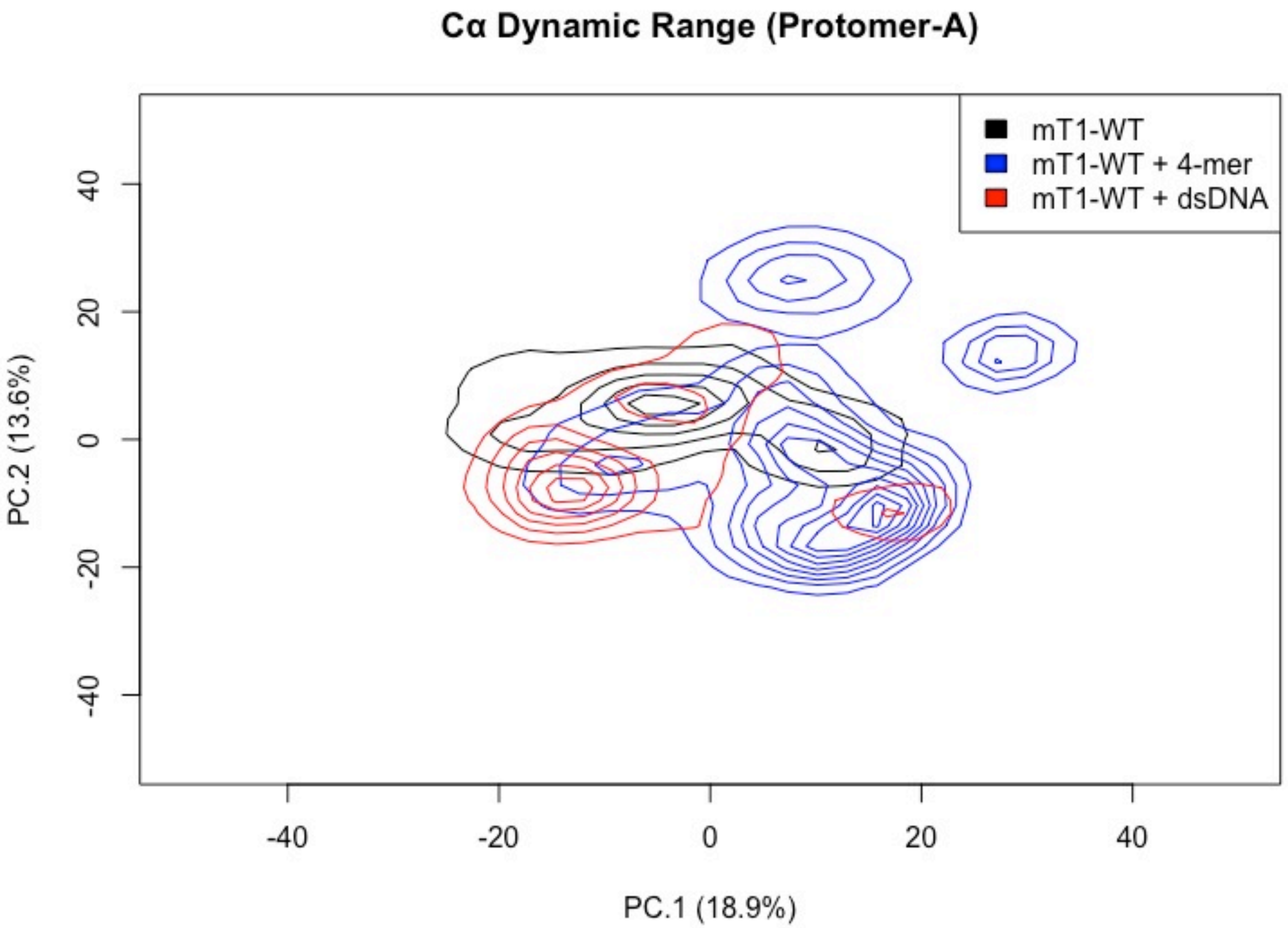

Cα Dynamic Range (Protomer-A)
mT1-WT
mT1-WT + 4-mer
mT1-WT + dsDNA
PC.2 (13.6%)
PC.1 (18.9%)
40
20
0
-20
-40
-40
-20
0
20
40

__Supplemental 4__: **Conformational free energy maps of mTREX1 backbone.** Cartesian coordinates of every Cα in mT1 were used to calculate map density functions as described in the relevant methods section. Density functions were plotted as contour maps, with innermost rings indicating the areas of highest density for respective systems. All calculations and graphing were performed in R v3.6.1.

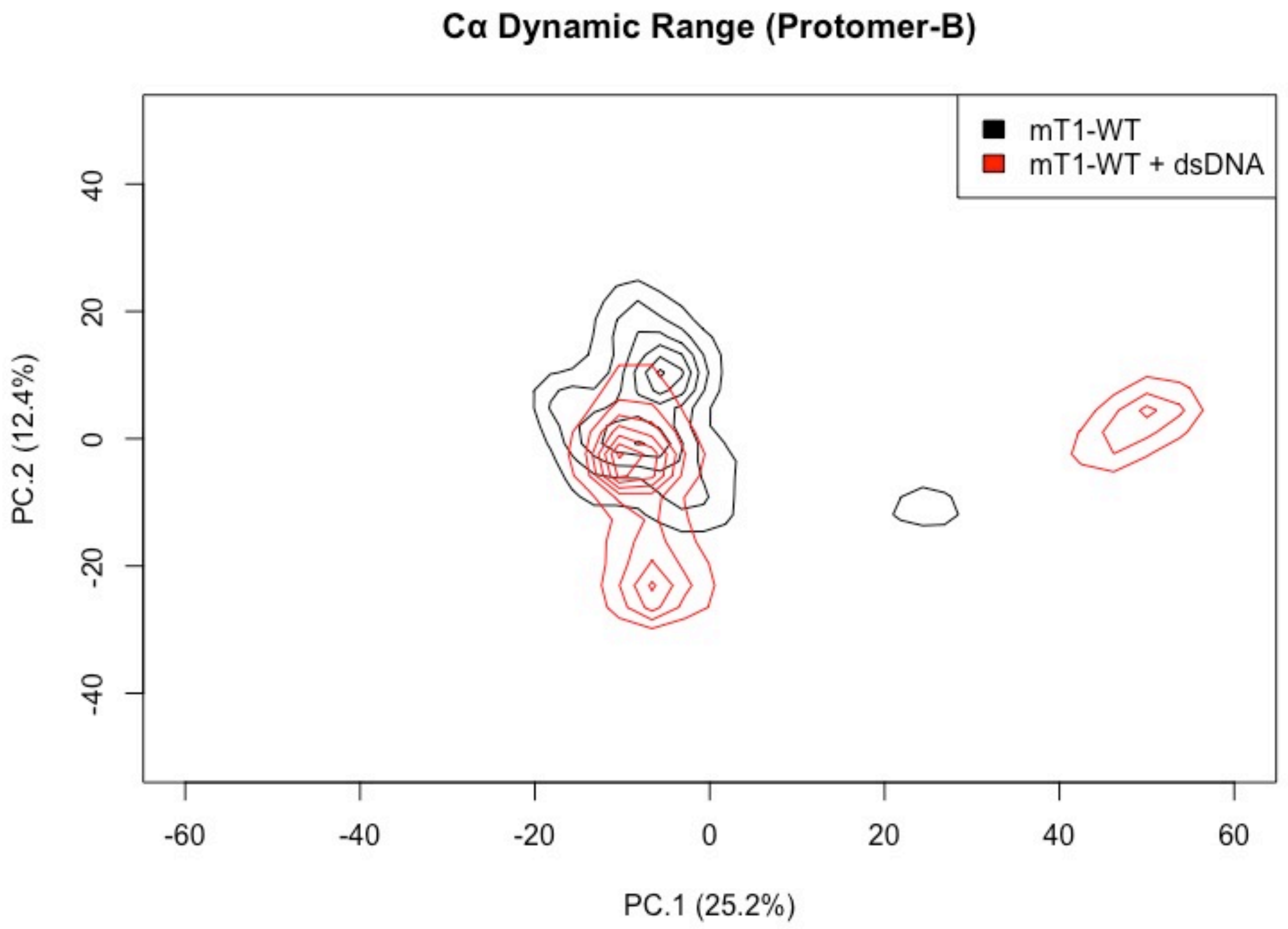
Cα Dynamic Range (Protomer-B)
mT1-WT
mT1-WT + dsDNA
PC.2 (12.4%)
PC.1 (25.2%)
-60
-40
-20
0
20
40
60
-40
-20
0
20
40

**Supplemental 5**: **Conformational free energy maps of mTREX1 backbone.** Cartesian coordinates of every Cα in the unbound protomer (‘Protomer-B’) of mT1 in every time frame in every simulation in both systems were used to calculate map density functions as described in the relevant methods section. Density functions were plotted as contour maps, with innermost rings indicating the areas of highest density for respective systems. All calculations and graphing were performed in R v3.6.1.

**Simulations Legend**

| Reaction | Description |
|---|---|
| *R1* | Supplemental 1A, ‘Ψ=0’ |
| *R2* | Supplemental 1A, ‘Ψ=0.25’ |
| *R3* | Supplemental 1A, ‘Ψ=0.5’ |
| *R4* | Supplemental 1A, ‘Ψ=0.75’ |
| *R5* | Supplemental 1A, ‘Ψ=1’ |
| *R6* | Supplemental 1B, ‘Ψ=0’ |
| *R7* | Supplemental 1B, ‘Ψ=0.25’ |
| *R8* | Supplemental 1B, ‘Ψ=0.5’ |
| *R9* | Supplemental 1B, ‘Ψ=0.75’ |
| *R10* | Supplemental 1B, ‘Ψ=1’ |
| *R11* | Supplemental 1C, ‘Ψ=0’ |
| *R12* | Supplemental 1C, ‘Ψ=0.25’ |
| *R13* | Supplemental 1C, ‘Ψ=0.5’ |
| *R14* | Supplemental 1C, ‘Ψ=0.75’ |
| *R15* | Supplemental 1C, ‘Ψ=1’ |
| *R16* | Supplemental 1D, ‘Ψ=0’ |
| *R17* | Supplemental 1D, ‘Ψ=0.25’ |
| *R18* | Supplemental 1D, ‘Ψ=0.5’ |
| *R19* | Supplemental 1D, ‘Ψ=0.75’ |
| *R20* | Supplemental 1D, ‘Ψ=1’ |
| *R21* | Supplemental 1E, ‘Ψ=0’ |
| *R22* | Supplemental 1E, ‘Ψ=0.25’ |
| *R23* | Supplemental 1E, ‘Ψ=0.5’ |
| *R24* | Supplemental 1E, ‘Ψ=0.75’ |
| *R25* | Supplemental 1E, ‘Ψ=1’ |
| *R26* | Figure 4A, red line |
| *R27* | Figure 4A, yellow line |
| *R28* | Figure 4A, green line |

| | |
|---|---|
| *R29* | Figure 4A, blue line |
| *R30* | Figure 4A, purple line |
| *R31* | Supplemental 2A, black line |
| *R32* | Supplemental 2A, red line |
| *R33* | Supplemental 2B, black line |
| *R34* | Supplemental 2B, red line |
| *R35* | Supplemental 2C, black line |
| *R36* | Supplemental 2C, red line |
| *R37* | Supplemental 2D, black line |
| *R38* | Supplemental 2D, red line |
| *R39* | Supplemental 2E, black line |
| *R40* | Supplemental 2E, red line |
| *R41* | Supplemental 2F, black line |
| *R42* | Supplemental 2F, red line |

### Simulations Parameters

| | **R1** | **R2** | **R3** | **R4** | **R5** | **R6** | **R7** | **R8** | **R9** | **R10** |
|---|---|---|---|---|---|---|---|---|---|---|
| $t$ (min) | 20 | 20 | 20 | 20 | 20 | 20 | 20 | 20 | 20 | 20 |
| $\Delta t$ (ms) | 1 | 1 | 1 | 1 | 1 | 1 | 1 | 1 | 1 | 1 |
| $[E]_0$ (M) | $10^{-12}$ | $10^{-12}$ | $10^{-12}$ | $10^{-12}$ | $10^{-12}$ | $10^{-12}$ | $10^{-12}$ | $10^{-12}$ | $10^{-12}$ | $10^{-12}$ |
| $[M]_0$ (M) | 0 | 0 | 0 | 0 | 0 | 0 | 0 | 0 | 0 | 0 |
| $\beta$ | 30 | 30 | 30 | 30 | 30 | 30 | 30 | 30 | 30 | 30 |
| $[S_\beta]_0$ (M) | 15e-9 | 15e-9 | 15e-9 | 15e-9 | 15e-9 | 15e-9 | 15e-9 | 15e-9 | 15e-9 | 15e-9 |
| $\Psi$ | 0 | 0.25 | 0.5 | 0.75 | 1 | 0 | 0.25 | 0.5 | 0.75 | 1 |
| $k_{-1}$ ($s^{-1}$) | 8.7 | 8.7 | 8.7 | 8.7 | 8.7 | 6.7 | 6.7 | 6.7 | 6.7 | 6.7 |
| $k_1$ ($M^{-1}s^{-1}$) | 2.1e8 | 2.1e8 | 2.1e8 | 2.1e8 | 2.1e8 | 1.6e8 | 1.6e8 | 1.6e8 | 1.6e8 | 1.6e8 |
| $k_2$ ($M^{-1}s^{-1}$) | 16 | 16 | 16 | 16 | 16 | 16 | 16 | 16 | 16 | 16 |
| $\Phi$ | Ψ | Ψ | Ψ | Ψ | Ψ | Ψ | Ψ | Ψ | Ψ | Ψ |
| $k_{-a}$ ($s^{-1}$) | $k_{-1}$ | $k_{-1}$ | $k_{-1}$ | $k_{-1}$ | $k_{-1}$ | $k_{-1}$ | $k_{-1}$ | $k_{-1}$ | $k_{-1}$ | $k_{-1}$ |
| $k_a$ ($M^{-1}s^{-1}$) | $k_1$ | $k_1$ | $k_1$ | $k_1$ | $k_1$ | $k_1$ | $k_1$ | $k_1$ | $k_1$ | $k_1$ |
| $k_b$ ($M^{-1}s^{-1}$) | 0 | 0 | 0 | 0 | 0 | 0 | 0 | 0 | 0 | 0 |

| | **R11** | **R12** | **R13** | **R14** | **R15** | **R16** | **R17** | **R18** | **R19** | **R20** |
|---|---|---|---|---|---|---|---|---|---|---|
| $t$ (min) | 20 | 20 | 20 | 20 | 20 | 20 | 20 | 20 | 20 | 20 |
| $\Delta t$ (ms) | 1 | 1 | 1 | 1 | 1 | 1 | 1 | 1 | 1 | 1 |
| $[E]_0$ (M) | $10^{-12}$ | $10^{-12}$ | $10^{-12}$ | $10^{-12}$ | $10^{-12}$ | $10^{-12}$ | $10^{-12}$ | $10^{-12}$ | $10^{-12}$ | $10^{-12}$ |
| $[M]_0$ (M) | 0 | 0 | 0 | 0 | 0 | 0 | 0 | 0 | 0 | 0 |
| $\beta$ | 30 | 30 | 30 | 30 | 30 | 30 | 30 | 30 | 30 | 30 |
| $[S_\beta]_0$ (M) | 15e-9 | 15e-9 | 15e-9 | 15e-9 | 15e-9 | 15e-9 | 15e-9 | 15e-9 | 15e-9 | 15e-9 |
| $\Psi$ | 0 | 0.25 | 0.5 | 0.75 | 1 | 0 | 0.25 | 0.5 | 0.75 | 1 |
| $k_{-1}$ ($s^{-1}$) | 4.5 | 4.5 | 4.5 | 4.5 | 4.5 | 2.2 | 2.2 | 2.2 | 2.2 | 2.2 |
| $k_1$ ($M^{-1}s^{-1}$) | 1.1e8 | 1.1e8 | 1.1e8 | 1.1e8 | 1.1e8 | 5.2e7 | 5.2e7 | 5.2e7 | 5.2e7 | 5.2e7 |

| | | | | | | | | | | |
|---|---|---|---|---|---|---|---|---|---|---|
| $k_2$ (M$^{-1}$s$^{-1}$) | 16 | 16 | 16 | 16 | 16 | 16 | 16 | 16 | 16 | 16 |
| $\Phi$ | Ψ | Ψ | Ψ | Ψ | Ψ | Ψ | Ψ | Ψ | Ψ | Ψ |
| $k_{-a}$ (s$^{-1}$) | $k_{-1}$ | $k_{-1}$ | $k_{-1}$ | $k_{-1}$ | $k_{-1}$ | $k_{-1}$ | $k_{-1}$ | $k_{-1}$ | $k_{-1}$ | $k_{-1}$ |
| $k_a$ (M$^{-1}$s$^{-1}$) | $k_1$ | $k_1$ | $k_1$ | $k_1$ | $k_1$ | $k_1$ | $k_1$ | $k_1$ | $k_1$ | $k_1$ |
| $k_b$ (M$^{-1}$s$^{-1}$) | 0 | 0 | 0 | 0 | 0 | 0 | 0 | 0 | 0 | 0 |

| | **R21** | **R22** | **R23** | **R24** | **R25** | **R26** | **R27** | **R28** | **R29** | **R30** |
|---|---|---|---|---|---|---|---|---|---|---|
| $t$ (min) | 20 | 20 | 20 | 20 | 20 | 20 | 20 | 20 | 20 | 20 |
| $\Delta t$ (ms) | 1 | 1 | 1 | 1 | 1 | 1 | 1 | 1 | 1 | 1 |
| $[E]_0$ (M) | $10^{-12}$ | $10^{-12}$ | $10^{-12}$ | $10^{-12}$ | $10^{-12}$ | $10^{-12}$ | $10^{-12}$ | $10^{-12}$ | $10^{-12}$ | $10^{-12}$ |
| $[M]_0$ (M) | 0 | 0 | 0 | 0 | 0 | 0 | 0 | 0 | 0 | 0 |
| $\beta$ | 30 | 30 | 30 | 30 | 30 | 30 | 30 | 30 | 30 | 30 |
| $[S_\beta]_0$ (M) | 15e-9 | 15e-9 | 15e-9 | 15e-9 | 15e-9 | 15e-9 | 15e-9 | 15e-9 | 15e-9 | 15e-9 |
| $\Psi$ | 0 | 0.25 | 0.5 | 0.75 | 1 | 0 | 0.25 | 0.5 | 0.75 | 1 |
| $k_{-1}$ (s$^{-1}$) | 0.25 | 0.25 | 0.25 | 0.25 | 0.25 | 8.7 | 6.7 | 4.5 | 2.2 | 0.25 |
| $k_1$ (M$^{-1}$s$^{-1}$) | 5.9e6 | 5.9e6 | 5.9e6 | 5.9e6 | 5.9e6 | 2.1e8 | 1.6e8 | 1.1e8 | 5.2e7 | 5.9e6 |
| $k_2$ (M$^{-1}$s$^{-1}$) | 16 | 16 | 16 | 16 | 16 | 16 | 16 | 16 | 16 | 16 |
| $\Phi$ | Ψ | Ψ | Ψ | Ψ | Ψ | Ψ | Ψ | Ψ | Ψ | Ψ |
| $k_{-a}$ (s$^{-1}$) | $k_{-1}$ | $k_{-1}$ | $k_{-1}$ | $k_{-1}$ | $k_{-1}$ | $k_{-1}$ | $k_{-1}$ | $k_{-1}$ | $k_{-1}$ | $k_{-1}$ |
| $k_a$ (M$^{-1}$s$^{-1}$) | $k_1$ | $k_1$ | $k_1$ | $k_1$ | $k_1$ | $k_1$ | $k_1$ | $k_1$ | $k_1$ | $k_1$ |
| $k_b$ (M$^{-1}$s$^{-1}$) | 0 | 0 | 0 | 0 | 0 | 0 | 0 | 0 | 0 | 0 |

| | **R31** | **R32** | **R33** | **R34** | **R35** | **R36** | **R37** | **R38** | **R39** | **R40** |
|---|---|---|---|---|---|---|---|---|---|---|
| $t$ (min) | 60 | 60 | 60 | 60 | 60 | 60 | 60 | 60 | 60 | 60 |
| $\Delta t$ (ms) | 1 | 1 | 1 | 1 | 1 | 1 | 1 | 1 | 1 | 1 |
| $[E]_0$ (M) | 7.5e-9 | 3.75e-9 | 7.5e-9 | 3.75e-9 | 7.5e-9 | 3.75e-9 | 7.5e-9 | 3.75e-9 | 7.5e-9 | 3.75e-9 |
| $[M]_0$ (M) | 0 | 3.75e-9 | 0 | 3.75e-9 | 0 | 3.75e-9 | 0 | 3.75e-9 | 0 | 3.75e-9 |
| $\beta$ | 1e4 | 1e4 | 1e4 | 1e4 | 1e4 | 1e4 | 1e4 | 1e4 | 1e4 | 1e4 |
| $[S_\beta]_0$ (M) | 0.83e-9 | 0.83e-9 | 0.83e-9 | 0.83e-9 | 0.83e-9 | 0.83e-9 | 0.83e-9 | 0.83e-9 | 0.83e-9 | 0.83e-9 |
| $\Psi$ | 0.75 | 0.75 | 0.80 | 0.80 | 0.85 | 0.85 | 0.90 | 0.90 | 0.95 | 0.95 |
| $k_{-1}$ (s$^{-1}$) | 0.10 | 0.10 | 0.050 | 0.050 | 0.020 | 0.020 | 0.010 | 0.010 | 0 | 0 |
| $k_1$ (M$^{-1}$s$^{-1}$) | 1e9 | 1e9 | 1e9 | 1e9 | 1e9 | 1e9 | 1e9 | 1e9 | 1e9 | 1e9 |
| $k_2$ (M$^{-1}$s$^{-1}$) | 10 | 10 | 10 | 10 | 10 | 10 | 10 | 10 | 10 | 10 |
| $\Phi$ | Ψ | Ψ | Ψ | Ψ | Ψ | Ψ | Ψ | Ψ | Ψ | Ψ |
| $k_{-a}$ (s$^{-1}$) | $k_{-1}$ | $k_{-1}$ | $k_{-1}$ | $k_{-1}$ | $k_{-1}$ | $k_{-1}$ | $k_{-1}$ | $k_{-1}$ | $k_{-1}$ | $k_{-1}$ |
| $k_a$ (M$^{-1}$s$^{-1}$) | $k_1$ | $k_1$ | $k_1$ | $k_1$ | $k_1$ | $k_1$ | $k_1$ | $k_1$ | $k_1$ | $k_1$ |
| $k_b$ (M$^{-1}$s$^{-1}$) | 0 | 0 | 0 | 0 | 0 | 0 | 0 | 0 | 0 | 0 |

| | **R41** | **R42** |
|---|---|---|
| $t$ (min) | 60 | 60 |
| $\Delta t$ (ms) | 1 | 1 |
| $[E]_0$ (M) | 7.5e-9 | 3.75e-9 |
| $[M]_0$ (M) | 0 | 3.75e-9 |
| $\beta$ | 1e4 | 1e4 |
| $[S_\beta]_0$ (M) | 0.83e-9 | 0.83e-9 |
| $\Psi$ | 1 | 1 |
| $k_{-1}$ (s$^{-1}$) | 0 | 0 |

| | | |
|---|---|---|
| $k_1$ ($M^{-1}s^{-1}$) | 1e9 | 1e9 |
| $k_2$ ($M^{-1}s^{-1}$) | 10 | 10 |
| $\Phi$ | Ψ | Ψ |
| $k_{-a}$ ($s^{-1}$) | $k_{-1}$ | $k_{-1}$ |
| $k_a$ ($M^{-1}s^{-1}$) | $k_1$ | $k_1$ |
| $k_b$ ($M^{-1}s^{-1}$) | 0 | 0 |

**Supplemental 6**: **Kinetic parameters for simulated reactions in these studies.** A table of the parameters used to simulate TREX1 exonuclease activity. A legend identifying the simulations is also provided. Tables were prepared in Word (Microsoft).
